# Evolutionary relationships and range evolution of greenhood orchids (subtribe Pterostylidinae): insights from plastid phylogenomics

**DOI:** 10.1101/2022.03.30.486312

**Authors:** Katharina Nargar, Kate O’Hara, Allison Mertin, Stephen Bent, Lars Nauheimer, Lalita Simpson, Heidi Zimmer, Bryan P.J. Molloy, Mark A. Clements

## Abstract

Australia harbours a rich and highly endemic orchid flora with over 90% of native species found nowhere else. However, little is known about the assembly and evolution of Australia’s orchid flora. Here, we used a phylogenomic approach to infer evolutionary relationships, divergence times, and range evolution in Pterostylidinae (Orchidoideae), the second largest subtribe in the Australian orchid flora, comprising the genera *Pterostylis* and *Achlydosa*. Phylogenetic analysis of 75 plastid genes provided well-resolved and supported phylogenies. Intrageneric relationships in *Pterostylis* were clarified and monophyly of eight of ten sections supported. *Achlydosa* was found to not form part of Pterostylidinae and instead merits recognition at subtribal level, as Achlydosinae.

Pterostylidinae were inferred to have originated in temperate eastern Australia in the early Oligocene, coinciding with the complete separation of Australia from Antarctica and the onset of the Antarctic Circumpolar Current, which led to profound changes in the world’s climate. Divergence of all major lineages occurred during the Miocene, accompanied by increased aridification and seasonality of the Australian continent, resulting in strong vegetational changes from rainforest to more open sclerophyllous vegetation. The majority of extant species were inferred to have originated in the Quaternary, from the Pleistocene onwards. The rapid climatic oscillations during the Pleistocene may have acted as important driver of speciation in Pterostylidinae. The subtribe underwent lineage diversification mainly within its ancestral range, in temperate eastern Australia. Long-distance dispersals to southwest Australia commenced from the late Miocene onwards, after the establishment of the Nullarbor Plain, which constitutes a strong edaphic barrier to mesic plants. Range expansions from the mesic into the arid zone of eastern Australia (Eremaean region) commenced from the early Pleistocene onwards. Extant distributions of Pterostylidinae in other Australasian regions, such as New Zealand and New Caledonia, are of more recent origin, resulting from long-distance dispersals from the Pliocene onwards. Temperate eastern Australia was identified as key source area for dispersals to other Australasian regions.

## 1 Introduction

Orchidaceae are the second largest angiosperm family with over 27,800 species and 750 genera (WFO, 2021; Chase et al., 2015). Since their origin in the Lower Cretaceous, ca. 112–137 Ma (Givnish et al., 2015; 2018; Serna-Sánchez et al., 2021; Silvestro et al., 2021), orchids have evolved a tremendous morphological and ecological diversity, including highly specialised mycorrhizal and plant-pollinator relationships (Dressler 1981; Pridgeon et al., 1999-2014). Orchidaceae are distributed worldwide, occur on all continents except Antarctica, and exhibit their highest species diversity in the tropics and subtropics (Pridgeon et al., 1999-2014).

Australia harbours a rich and highly endemic orchid flora of more than 1,600 species, with over 90% of Australia’s native orchids endemic to the country (Jones, 2021). The Australian orchid flora is especially rich in terrestrial orchids from subfamily Orchidoideae, harbouring over one third of the global diversity of this subfamily (WCSP, 2018). For several lineages within Orchidoideae, such as subtribes Pterostylidinae, Caladeniinae, Diuridinae, and Prasophyllinae, the centre of diversity lies in Australia (Pridgeon et al., 1999-2014). However, the spatio-temporal evolution of many Australasian orchid lineages is still poorly understood (Givnish et al., 2016; Nauheimer et al., 2018; Nargar et al., 2018).

Pterostylidinae constitutes the second largest subtribe in the Australian orchid flora, with over 300 species (Jones, 2021). In its traditional circumscription, the subtribe comprises one genus, *Pterostylis* R.Br. (Dressler, 1993). Pterostylidinae are geophytic herbs with root-stem tuberoids, characterised by flowers with a hoodlike structure (termed galea) formed by the median sepal and lateral petals, partially fused lateral sepals forming the synsepalum, and an irritable mobile labellum (Figure 1) (Jones and Clements 2002a; Pridgeon et al., 2003). Pterostylidinae are predominantly pollinated by fungus gnats of the families Mycetophilidae, Phoridae, and Culicidae (order Diptera) which become temporarily trapped in the hood-shaped flowers to facilitate pollination (Pridgeon et al., 2003; Kuiter, 2015).

**Figure 1.**
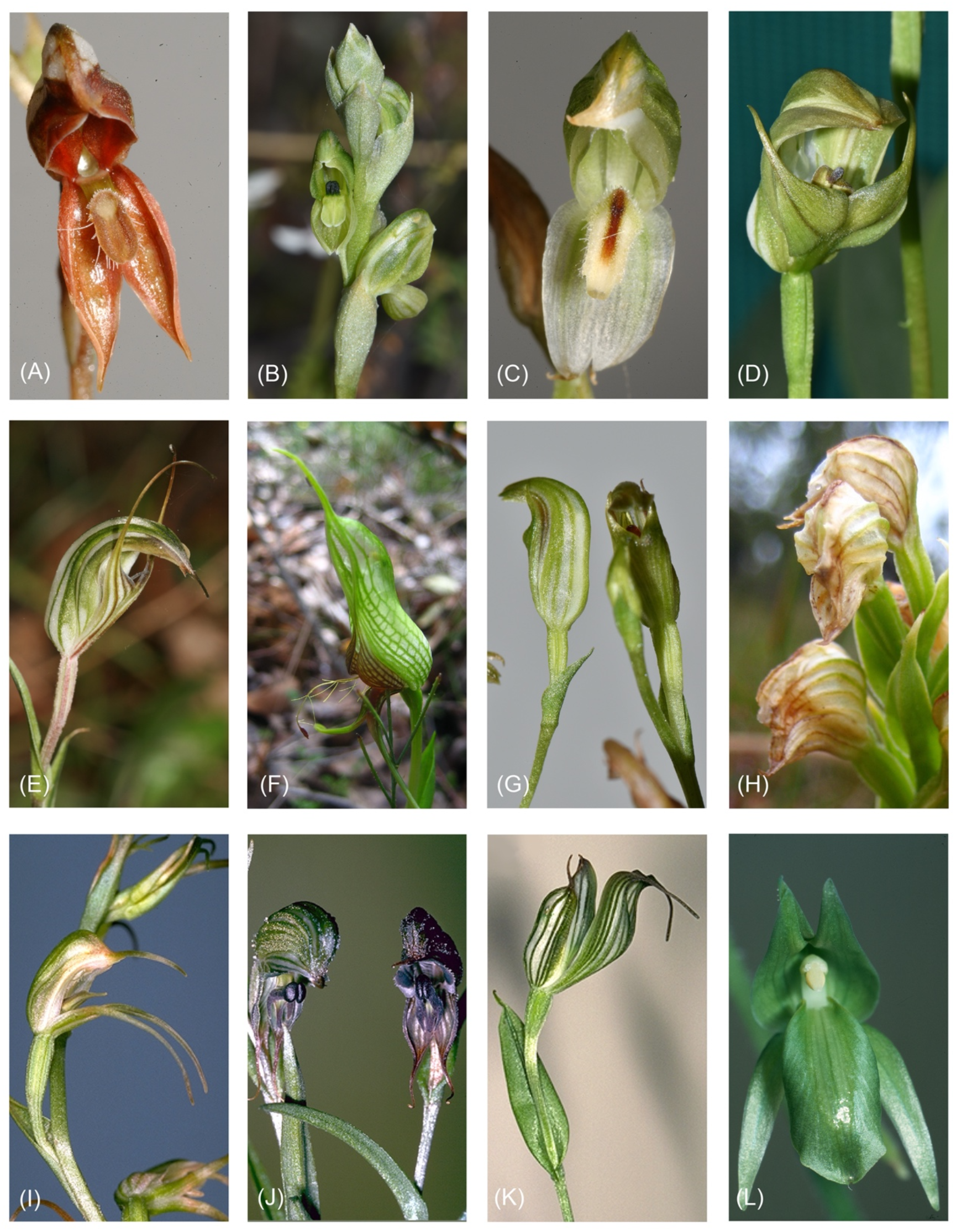
Morphological diversity within Pterostylidinae. (A) *Pterostylis rufa* (sect. *Oligochaetochilus*); (B) *Pterostylis bicolor* (sect. *Hymenochilus*); (C) *Pterostylis longifolia* (sect. *Squamatae*); (D) *Pterostylis curta* (sect. *Pterostylis*); (E) *Pterostylis striata* (sect. *Foliosae);* (F) *Pterostylis barbata* (sect. *Catochilus*); (G) *Pterostylis parviflora* (sect. *Parviflorae*); (H) *Pterostylis daintreeana* (sect. *Pharochilum*); (I) *Pterostylis sargentii* (sect. *Urochilus*); (J) *Pterostylis recurva* (sect. *Stamnorchis*), and (K) *Achlydosa glandulosa*.

The subtribe has a predominantly Australasian distribution with centre of diversity in Australia (289 sp.; Jones 2021), extending to New Zealand (16 sp.; Breitwieser et al., 2010), New Caledonia (7 sp.; Endemia, 2021), Indonesia (3 sp.; Schuiteman et al., 2008), Papua New Guinea (2 sp.; de Vogel et al., 2021), and East Timor (1 sp., Silveira et al., 2008). Pterostylidinae are primarily found in mesic habitats, from near sea level to ca. 3500 m (de Vogel et al., 2021). In Australia, Pterostylidinae are most diverse in the mesic zone of temperate southeast Australia (Jones, 2021; ALA, 2021). A family-wide phylogenetic study inferred an Australian/Pacific or Australian origin of Pterostylidinae in the Eocene, ca. 38.2 Ma (Givnish et al., 2016). However, little is known about range evolution of Pterostylidinae through time.

Before the molecular phylogenetics era, subtribe Pterostylidinae was placed in tribe Diurideae (Dressler, 1993). However, molecular phylogenetic studies demonstrated that Pterostylidinae belonged to tribe Cranichideae, the sister group to Diurideae (Cameron et al., 1999; Kores et al., 2001; Clements et al., 2002; Givnish et al., 2015; Serna-Sánchez et al., 2021; Perez-Escobar et al., 2021). The concept of Pterostylidinae was expanded by Chase et al. (2015) to include New Caledonian monotypic genus *Achlydosa* M.A.Clem. & D.L.Jones based on its phylogenetic proximity and similarities in floral morphology of its sole species, *Achlydosa glandulosa* (Schltr.) M.A.Clem. & D.L.Jones. However, a sister group relationship between *Pterostylis* and *Achlydosa* and thus the monophyly of Pterostylidinae *sensu* Chase et al. (2015) has not been firmly established as phylogenetic studies resulted in different topologies within early diverging Cranichideae (Gustavson et al., 2010; Cisternas et al., 2012; Gamisch et al., 2015; Givnish et al., 2015).

To accommodate the morphological diversity in Pterostylidinae, different classifications have been proposed over the past two centuries (Brown, 1810; Don, 1830; Reichenbach, 1871; Bentham, 1873; Rupp, 1933; Szlachetko, 2001; Jones and Clements, 2002b; Janes and Duretto, 2010; Chase et al., 2015; Jones, 2015; Clements and Jones, 2016; Jones and Clements, 2017). While for a long time only a single genus, *Pterostylis*, was recognised within the subtribe, Szlachetko (2001) proposed the segregation of *Pterostylis* into three genera by erecting *Oligochaetochilus* Szlach. and *Plumatichilos* Szlach., which resulted in non-monophyletic taxa as shown in subsequent phylogenetic analysis based on ITS data (Clements et al. 2011; Figure 2). Based on morphological studies, Jones and Clements (2002a) initially distinguished 12 informal groups within *Pterostylis*. Based on a combined analysis of morphological characters and ITS data (Jones and Clements, 2002a), Jones and Clements (2002b) further divided Pterostylidinae with the aim to render taxonomic groups within Pterostylidinae monophyletic, resulting in a total of 16 genera (Figure 2).

**Figure 2.**
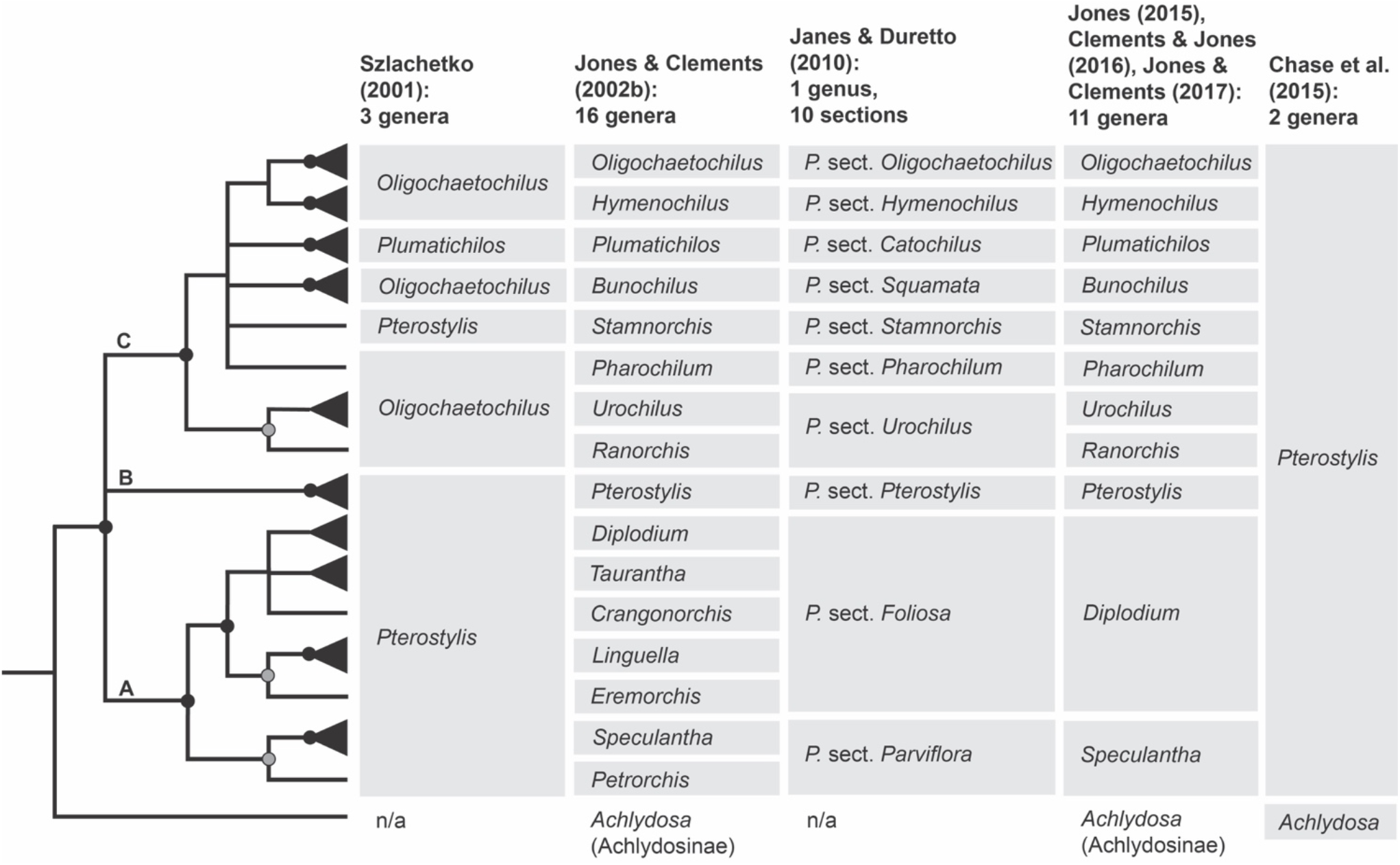
Summary cladogram of phylogenetic relationships in Pterostylidinae based on Clements et al. (2011; modified) and systematic concepts in Pterostylidinae over the past two decades. Solid black circles denote nodes which received maximum support (Bayesian posterior probabilities of 1); grey circles denote moderately supported notes (posterior probabilities of 0.98-0.99). Monotypic taxa are depicted with a single line as terminals. Branches with lower support values (<0.95) are collapsed. Grey boxes highlight taxa which are considered part of Pterostylidinae in the respective classification. Taxa described prior to 2001 and publication years: sect. *Catochilus* Benth. (1873); *Diplodium* Sw. (1810); sect. *Foliosa* G.Don (1830); sect. *Parviflora* Benth. (1873); *Pterostylis* R.Br. (1810); sect. *Squamata* G.Don (1830). For historic infrageneric classification systems for Pterostylidinae, see Jones and Clements (2002a).

To assess phylogenetic support of the taxonomic groups *sensu* Jones and Clements (2002b), Janes et al. (2010) reanalysed published ITS data from Pterostylidinae with 12 additional *Pterostylis* species. Janes et al. (2010) found high support for three main lineages within Pterostylidinae, termed clades A, B and C, however relationships among these three lineages remained unclear due to low nodal support.

Janes and Duretto (2010) presented a revised classification for Pterostylidinae in which all 15 segregate genera *sensu* Jones and Clements (2002b) were sunk into *Pterostylis s.l*.. The infrageneric classification of Janes and Duretto (2010) was based on a combination of phylogenetically supported lineages and/or morphological evidence, and partly aligned with taxonomic delineations of Jones and Clements (2002b): seven sections directly corresponded to taxonomic groupings *sensu* Jones and Clements (2002b) (Figure 2). In two instances, taxonomic concepts were broadened to accommodate two taxa *sensu* Jones and Clements (2002b) and sinking these to sectional level: *Ranorchis* and *Urochilus* were sunk into sect. *Urochilus*, and *Petrorchis* and *Speculantha* were sunk into sect. *Parviflorae* (Figure 2). Further, the five genera *Crangonorchis, Diplodium, Eremorchis, Linguella*, and *Taurantha*, were sunk into sect. *Foliosae* due to lack of resolution among these taxa (Figure 2). Given the limited resolution and support of the phylogenetic inferences based on ITS data, Janes and Duretto (2010) stated that additional study was required to determine whether further revisions are warranted.

Clements et al. (2011) presented the most comprehensively sampled phylogenetic study of Pterostylidinae to date, comprising 152 species, based on nuclear data (ITS) and for a subsample the addition of one plastid marker (*mat*K). While taxon sampling was more extensive, many of the previously unresolved relationships among major lineages remained unclear. Subsequently, Jones and Clements (Jones 2015, 2021; Clements and Jones, 2016; Jones and Clements, 2017) revised the generic classification of Pterostylidinae to recognise the more broadly defined sect. *Parviflorae* and sect. *Foliosae* (Janes and Duretto, 2010) at generic level (as *Speculantha s.l.* and *Diplodium s.l*.; Figure 2). The Council Heads of Australasian Herbaria (CHAH, 2018) recommended the use of the broader taxonomic circumscription of *Pterostylis*, which was adopted in the Australian Plant Census and in most Australasian herbaria. However, the more narrowly defined treatment of Pterostylidinae (Jones and Clements, 2002b; Jones, 2015; Clements and Jones, 2016; Jones and Clements, 2017) has remained in use as alternative classification (e.g., Jones, 2006; Jones, 2021). The use of a dual taxonomic classification system for Pterostylidinae has resulted in confusion and inconsistencies in the use of taxonomic names, e.g., in online biodiversity databases such as the Australasian Virtual Herbarium (AVH, 2022) and Atlas of Living Australia (ALA, 2022).

As previous efforts to assess the monophyly of taxa within Pterostylidinae in a phylogenetic framework were hampered by limited resolution and support of inferred evolutionary relationships (Jones and Clements, 2002a; Janes et al., 2010; Clements et al., 2011) (Figure 2), further molecular study is required to resolve phylogenetic relationships within the subtribe. Lack of resolution of evolutionary relationships in Pterostylidinae also precluded understanding of range evolution of this diverse Australasian orchid lineage in a temporal framework.

This study aims to clarify evolutionary relationships within Pterostylidinae based on plastid phylogenomics, to infer divergence times and range evolution of the subtribe to understand its biogeographic history in the context of paleogeographic and paleoclimatic changes, and to provide a robust phylogenetic framework to allow for a re-evaluation of taxonomic concepts in Pterostylidinae.

## 2 Materials and Methods

### 2.1 Taxon sampling

In total, 150 orchid samples were included in the study. For Pterostylidinae, 98 species (106 accessions) were sampled. As outgroups, a total of 43 species (44 accessions) representing all five subfamilies of Orchidaceae were included. Within Orchidoideae, representatives of all four tribes (Codonorchideae, Cranichideae, Diurideae, and Orchideae) were sampled and Epidendroideae were represented by 14 samples from six Epidendroideae tribes. For 18 outgroup samples, plastid and nuclear data from previous molecular studies were sourced from GenBank. Sample details are provided in Supplementary Material S1. Taxonomic concepts for Pterostylidinae at genus-level follow the recommendations of the Council Heads of Australasian Herbaria (CHAH, 2018). Corresponding synonyms for the classification of Pterostylidinae *sensu* Jones and Clements (2002b) including subsequent revisions (Jones, 2015; Clements and Jones, 2016; Jones and Clements, 2017) are provided in Supplementary Material S1.

### 2.2 DNA isolation, library construction, and sequencing

For DNA extractions, 10-20 mg of silica-dried stem or leaf tissue was ground with a Qiagen TissueLyser II (Qiagen, Melbourne, Vic., Australia). DNA was extracted using the DNeasy 96 Plant Kit or DNeasyPlant Mini Kit (Qiagen) following the manufacturer’s protocol and eluted in 100 μL of TE buffer (Qiagen).

Sequencing libraries were constructed from 100 ng of total DNA using the TruSeq Nano DNA LT library preparation kit (Illumina, San Diego, CA, USA) for an insert size of 350 base pairs (bp), following the manufacturer’s protocol. Sequencing libraries were multiplexed 96 times and DNA sequencing with 125-bp paired-end reads was performed on an Illumina HiSeq 2500 platform at the Australian Genomic Research Facility (AGRF, Melbourne, Vic., Australia).

### 2.3 DNA sequence assemblies and alignments

Raw sequences were trimmed applying a Phred score > 20 using Trimmomatic 0.39 (Bolgner et al., 2014), and deduplicated using clumpify from BBtools 38.9 (Bushnell, 2014). Read pairs were then assembled using SPAdes 3.15 (Bankevich et al., 2012). Plastid and nuclear ribosomal rRNA databases were extracted from NCBI’s Nucleotide Entrez database using Entrez Programming Utilities (2008) using taxonomic, keyword, and sequence length constraints. Contigs were identified as derived from plastid or nuclear ribosomal rRNA source using blastn against these extracted databases. Genes within plastid and nuclear ribosomal rRNA contigs were identified by homology using BLAST (Altschul et al., 1990) and BLASTx (RRID:SCR_001653) against genes extracted from annotations of the reference sequence sets extracted from nuccore. In cases where *de novo* assemblies showed evidence of misassembled regions, reference-guided assemblies were carried out with a reference sequence from a closely related species using Geneious Prime 2020.0.1 (Biomatters Ltd., https://www.geneious.com).

DNA sequences for each locus were aligned separately using MAFFT multiple alignment software (ver. 7.388, see https://mafft.cbrc.jp/alignment/software/; Katoh and Standley, 2013) as implemented in Geneious Prime 2020.0.1. Two alignments were generated: a plastid dataset comprising 75 genes (87 exons and 12 introns) and a nuclear dataset of the ribosomal rRNA cistron including the external and internal transcribed spacers (ETS, ITS). We refrained from combining the two datasets because of the strong imbalance between plastid versus nuclear loci which has the potential to drown out moderate phylogenetic signal from the smaller nuclear partition. For coding regions, start and stop codons were visually verified in Geneious Prime 2020.0.1. Sequences featuring frame shift-inducing mutations and resulting internal stop codons were excluded from final alignments and subsequent analyses. The plastid alignment totalled 91,090 bp in length and the nuclear alignment 8,808 bp. Both datasets were partitioned into coding and non-coding regions and this partitioning was applied in subsequent phylogenetic analyses.

### 2.4 Phylogenomic analyses

Phylogenetic relationships were reconstructed using both maximum likelihood (ML) and Bayesian inference (BI) with the best-fit substitution model applied as determined for each partition. ML analyses were conducted in both IQ-TREE 1.6.1 (Nguyen et al., 2015) and RAxML 8.2.4 (Stamatakis, 2014). Model selection in IQ-TREE was performed based on the Bayesian information criterion (BIC; Schwarz, 1978) using ModelFinder (Kalyaanamoorthy et al., 2017). For the plastid dataset, the GTR+F+R5 model was identified as the best-fit model for the protein-coding genes and the K3Pu+F+R2 model for the intronic partition. For the nuclear dataset, the GTR+F+R4 and GTR+F+I+G4 models were selected for the rRNA and spacer partitions, respectively. The chosen models were incorporated into subsequent ML analyses in IQ-TREE which were performed separately for each dataset using the edge-proportional partition model (Chernomor et al., 2016). Bootstrap support (BS) values were calculated under the same models, applying the ultrafast bootstrap algorithm (UFBoot; Hoang et al., 2018) for 1000 pseudo replicates. Model selection for ML analysis in RAxML and MrBayes 3.2.7a (Ronquist and Huelsenbeck, 2003) was performed based on the Akaike information criterion (AIC) using MrModeltest 2.4 (Nylander, 2004) in PAUP* 4.0a (Swofford, 2003). The GTR+I+G model was identified as the best-fit model of nucleotide evolution for all data partitions. It was incorporated into subsequent partitioned ML analyses in RAxML with the rapid bootstrap algorithm in effect for 1000 pseudo replicates. The same data partitioning and substitution model was implemented for BI conducted in MrBayes 3.2.7a (Ronquist and Huelsenbeck, 2003) on XSEDE via the CIPRES Science Gateway (Miller et al., 2010; https://www.phylo.org). Three independent Markov-chain Monte Carlo (MCMC) searches were run per dataset with four chains of five million generations, ensuring the standard deviation of split frequencies was below 0.01. Trees were sampled at a frequency of 500 generations and a burn-in fraction of 20% was discarded. Majority-rule consensus trees including posterior probabilities (PP) were generated from the post burn-in sample.

### 2.5 Divergence-time estimation

Bayesian divergence-time estimation was carried out in BEAST 2.6.1 (Bouckaert et al., 2019) on XSEDE via the CIPRES Science Gateway. A secondary calibration approach was adopted for the molecular clock analysis due to an absence of fossil records in Pterostylidinae. Estimated node ages, including 95% highest posterior density estimates (HPD), were taken from a plastid phylogenomic study in monocots (Givnish et al., 2018). The plastid dataset was reduced to the 25 most parsimony informative plastid coding regions (44,510 bp) as identified using PAUP* v.4.0a to address computational limitations. The plastid and nuclear alignments were analysed separately.

For analysis of the plastid dataset, four normally distributed priors were set based on Givnish et al. (2018): at the Vanilloideae stem node (offset = 72.9 Ma, SD = 2.7), the Cypripedioideae stem node (offset = 64.2 Ma, SD = 2.6), the Epidendroideae stem node (offset = 53.6 Ma, SD = 2.0) and the Orchidoideae crown node (offset = 44.9 Ma, SD = 1.9). For the nuclear dataset, secondary calibrations were set as normally distributed priors for two nodes: the Orchidoideae crown node (offset = 44.9 Ma, SD = 1.9) and the Cranichideae stem node (offset = 40.2 Ma, SD = 1.9). For the plastid dataset, the backbone tree topology was constrained according to Givnish et al. (2018) with monophyly enforced at the Vanilloideae and Cypripedioideae stem nodes and for the nuclear dataset at the Orchidoideae crown node and Cranichideae stem node.

Divergence-time estimation was carried out using both a strict clock and an uncorrelated relaxed lognormal clock with both pure-birth (Yule) and birth-death models selected for the speciation/extinction process. The previously selected GTR+I+G substitution model was in effect with four gamma categories and empirical base frequencies. For each dataset, ten analyses were performed under the strict clock model, each with ten million generations and sampling every 1,000 generations. Under the relaxed clock model, multiple runs were performed (>15) for each dataset with 100 million generations and sampling every 10,000 generations. For each clock model, a single analysis with an empty dataset was conducted to evaluate the influence of the selected priors on the resulting posteriors. The MCMC trace files were visualised in Tracer 1.7.1 (Rambaut et al., 2018), assessing effective sample sizes of parameters and burn-in fraction. Sampled trees from each run were combined in LogCombiner 2.6.1 (Drummond and Rambaut, 2007), excluding a burn-in of 10% and resampling of 10,000 trees. Maximum-clade credibility chronograms including mean node heights and 95% HPD values were generated in TreeAnnotator 2.6.0 (Drummond and Rambaut, 2007). The various clock and speciation models were evaluated using a posterior simulation-based analogue of Akaike’s information criterion (AIC; Akaike 1974), termed AICM (Raftery et al., 2007), as implemented in AICM Analyser included in BEAST 2.6.1.

### 2.6 Ancestral range estimation

Within continental Australia, the delineation of biogeographic subregions was based on the terrestrial phytoregionalisation defined by Ebach et al. (2015) and slightly modified to reflect distribution patterns in Pterostylidinae. The following seven biogeographic areas were coded: a: Euronotian region; b: southwest Australia; c: Eremaean region (limited to inland eastern Australia); d: northern region (sub-region Atherton); e: Lord Howe Island; f: New Zealand, and g: New Caledonia. Distributions were sourced from Breitwieser et al. (2010), Jones (2021), and Endemia (2021). Distributions in Indonesia, Papua New Guinea, and Timor and were not included in the area coding as *Pterostylis* species with distributions in these regions were not available for this study.

Ancestral-range estimations were conducted using the R package BioGeoBEARS (see https://github.com/nmatzke/BioGeoBEARS; Matzke, 2013), based on the plastid maximum clade-credibility chronogram from the BEAST divergence dating analysis, pruned of duplicate samples of species and of all outgroups to Pterostylidinae *s.s.*. We refrained from an ancestral-range estimations based on the nuclear dataset because relationship among several major lineages within Pterostylidinae were poorly supported and resolution was low in several terminal clades. For the BioGeoBEARS analysis based on the plastid dataset, we implemented three models of biogeographic range inheritance: dispersal-extinction-cladogenesis (DEC; Ree and Smith, 2008), a ML version of Ronquist’s parsimony dispersal-vicariance (DIVA; Ronquist, 1997), termed DIVALIKE (Matzke, 2013), and a simplified likelihood interpretation of the Bayesian “BayArea” program (Landis et al., 2013) known as BAYAREALIKE (Matzke, 2013). We refrained from including jump dispersal (+J) due to conceptual and statistical problems identified by Ree and Sanmartin (2018). The maximum number of combined areas was set to the maximum number of observed areas in species (5), and equal probabilities were applied to all dispersal events. The likelihood values were compared using AIC, and the best-fit model was used to infer the relative probabilities of ancestral ranges at each node in the phylogeny.

## 3 Results

### 3.1 Phylogenomic analysis

Phylogenetic reconstructions based on 75 plastid genes based on maximum likelihood with IQ-TREE, RAxML, and Bayesian inference yielded congruent results (Figure 3, Figure 4, Supplementary Material S2.1, S2.2). In the following, results of the IQ-TREE analysis are presented.

**Figure 3:**
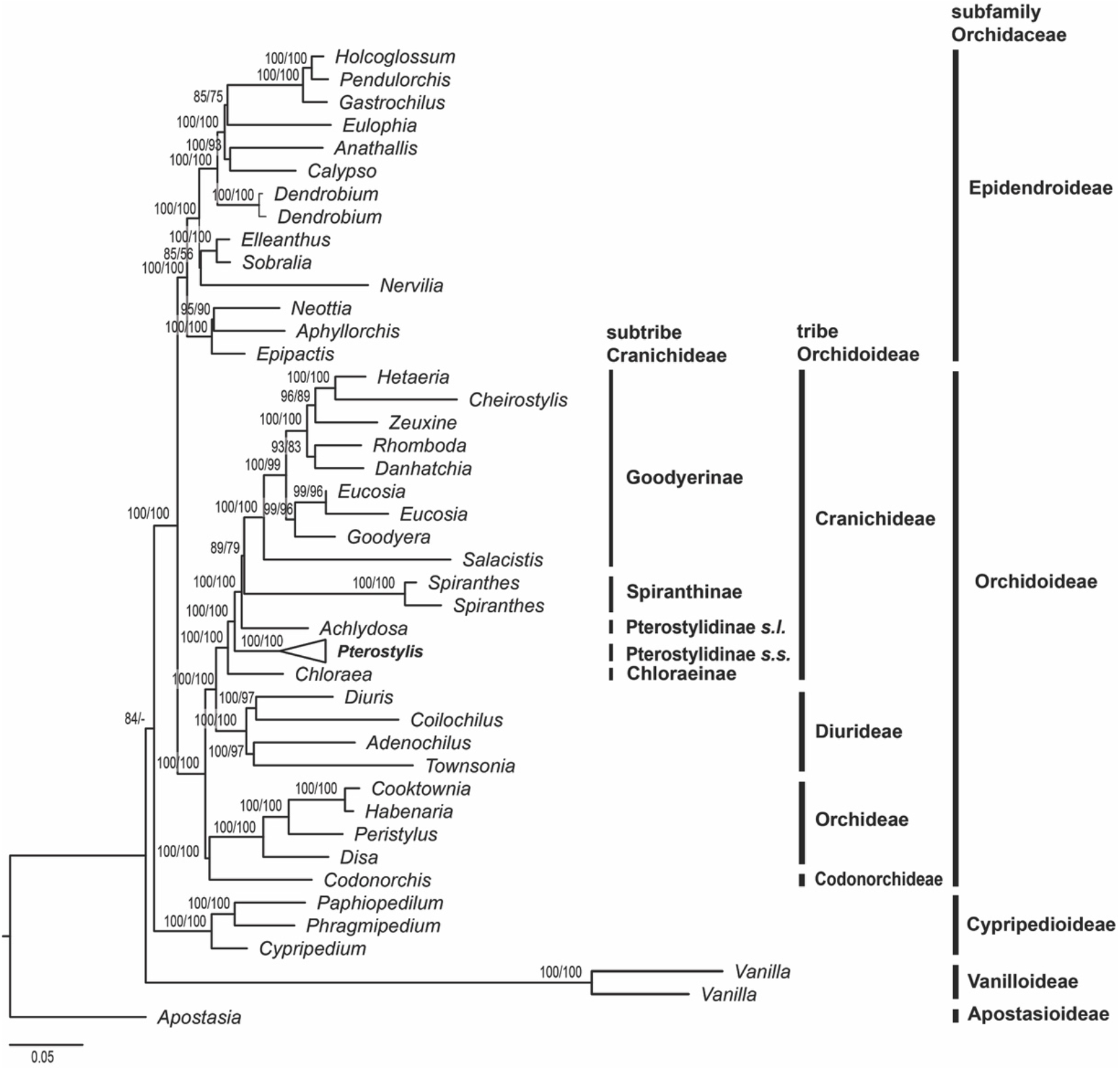
Higher-level phylogenetic relationships in Orchidaceae and placement of Pterostylidinae within Cranichideae. Maximum likelihood reconstruction based on 75 plastid genes (91,090 bp alignment) with IQ-TREE Nodal support values > 50 are given above branches (ultrafast bootstrap values from IQ-TREE analysis followed by bootstrap values from RAxML analysis).

**Figure 4:**
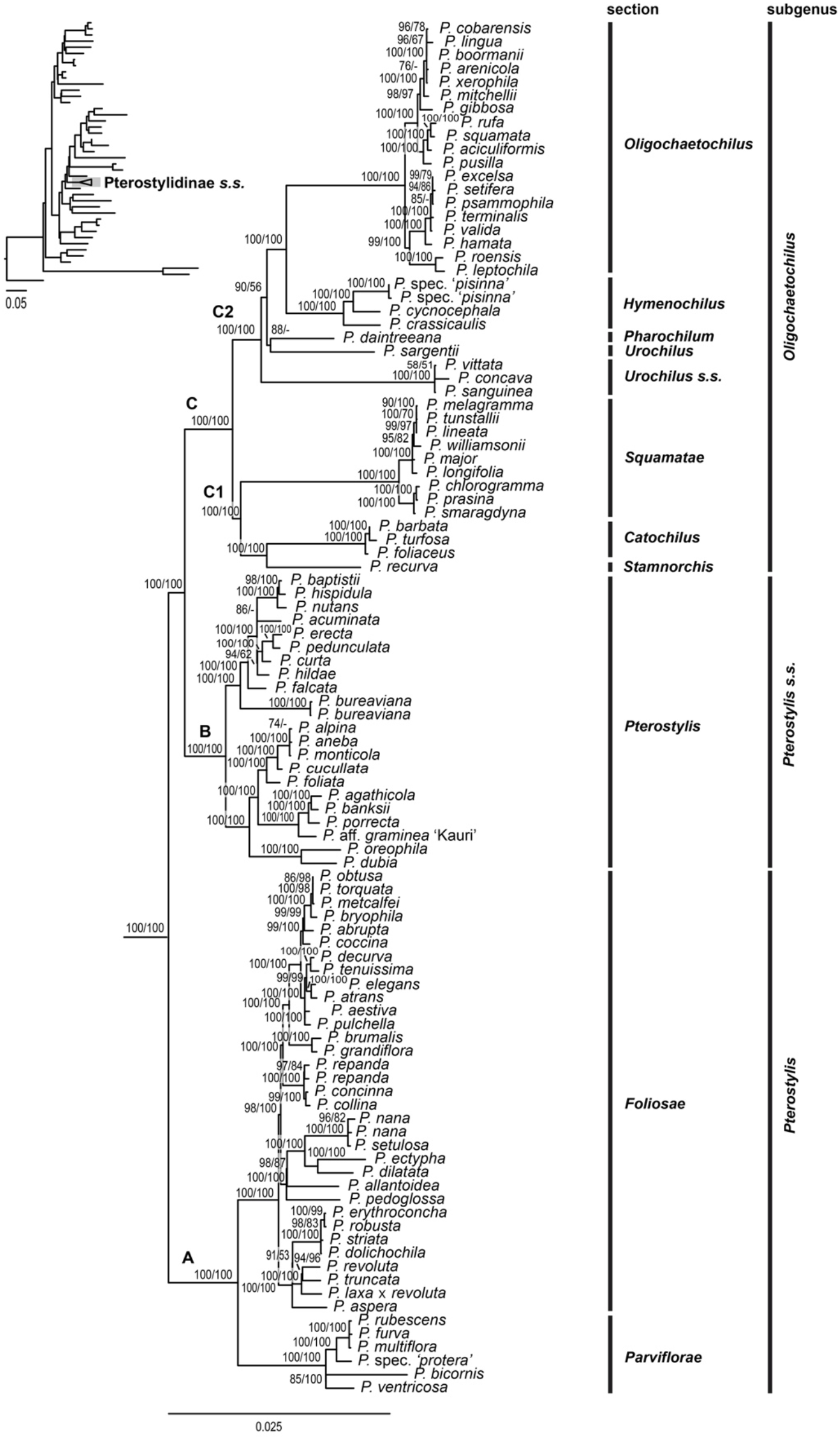
Phylogenetic relationships within *Pterostylis* based on maximum likelihood analysis of 75 plastid genes (91,090 bp alignment) with IQ-TREE. Labels A, B, and C refer to the three major clades within the genus, C1 and C2 represent the two main clades within clade C. Nodal support values > 50 are given above branches (ultrafast bootstrap values from IQ-TREE analysis followed by bootstrap values from RAxML analysis). Tree insert shows phylogenetic position of Pterostylidinae in orchid phylogeny (see Figure 3).

#### Higher-level phylogenetic relationships in Orchidaceae and phylogenetic placement of Pterostylidinae within Cranichideae

The phylogenetic reconstruction based on the 75 plastid genes showed subfamily Apostasiodeae as the first-diverging lineage, followed by Vanilloideae, Cypripedioideae, Orchidoideae, and Epidendroideae (Figure 3). Within subfamily Orchidoideae, tribe Codonorchideae was resolved as sister group to Orchideae, and the two tribes in turn were sister group to a clade comprising Diurideae as sister group to Cranichideae. The phylogenetic relationships among the four tribes within subfamily Orchidoideae received maximum support (Figure 3). Within tribe Cranichideae, subtribe Chloraeinae diverged first, followed by *Pterostylis* (Pterostylidinae *s.s.*). Next diverging was a clade comprised of *Achlydosa* (Pterostylidinae *s.l.*) in sister group position to the remainder of Cranichideae. These relationships received maximum support values (Figure 3).

#### Phylogenetic relationships within *Pterostylis*

Within *Pterostylis*, monophyly of the three major lineages, clades A, B, and C, received maximum support (Figure 4). Clade A was sister group to clade B and clade C with maximum support. Clade A comprised sections *Foliosae* and *Parviflorae*, both receiving maximum support, clade B harboured sect. *Pterostylis*, and clade C the remaining seven sections.

Within clade C, two main lineages were resolved with maximum support, termed clades C1 and C2 here. In clade C1, sect. *Stamnorchis* was sister group to a highly supported sect. *Catochilus* and the two in turn were sister group to a highly supported sect. *Squamatae.* These relationships received maximum support (Figure 4).

Within clade C2, relationships among sect. *Urochilus s.s., P. sargentii* (part of sect. *Urochilus s.l.*), and *P. daintreeana* (sect. *Pharochilum*) remained unclear due to differing topologies in the ML analyses. The reconstruction with IQ-TREE showed *Urochilus s.s.* as first diverging lineage, followed by a clade with *P. daintreeana* and *P. sargentii*, sister to the remainder of clade C2, whereas the reconstruction with RAxML showed a basal grade with *P. sargentii* diverging first, followed by sect. *Urochilus s.s.*, and *P. daintreeana*, sister to the remainder of clade C2. These relationships were not or only weakly-moderately supported. The next diverging lineage within clade C2 received maximum support and harboured a highly supported sect. *Hymenochilus* as sister group to a highly supported sect. *Oligochaetochilus* (Figure 4).

#### Phylogenetic relationships – nuclear data

Phylogenetic analysis of the nuclear data yielded congruent results to those of the plastid dataset. Overall, resolution and/or nodal support values were often lower in the nuclear analysis.

Relationships among several of the main lineages remained poorly supported, e.g., relationships within subgenus *Oligochaetochilus*. Relationships at among closely related species often remained poorly resolved (e.g., within sections *Foliosae, Oligochaetochilus*, and *Squamata*). (Supplementary Material S2.3, S2.4, S2.5, S 2.6).

Within *Pterostylis*, the three main clades, A, B, and C, were highly supported. Clade A was retrieved in sister group position to clade B, and the two in turn were found as sister group to clade C, however the sister group relationship between clades A and B received only weak support. Within clade A, the monophyly of sect. *Parviflorae* and of sect. *Foliosae* was highly supported as well as their sister group relationship. Within clade C, sections *Catochilus*, *Hymenochilus, Oligochaetochilus*, and *Squamatae* were highly supported. However, phylogenetic relationships among the main lineages within clade C remained largely unclear due to lack of nodal support. The only highly supported relationships among major lineages within the clade was the sister group relationship between sections *Hymenochilus* and *Oligochaetochilus*, in congruence with the results based on the plastid dataset (Supplementary Material S2.4).

### 3.2 Divergence time estimation

The evaluation of divergence time analyses based on different combinations of molecular clock and speciation models using AICM (Raftery et al. 2007) determined the relaxed clock model as best fit model for the plastid and the nuclear dataset (Supplementary Material S3.1). As best-fit speciation models, the birth-death process was determined for the plastid dataset and the Yule process for the nuclear dataset (Supplementary Material S3.1).

#### Divergence time estimations based on plastid dataset

Tribes Cranichideae and Diurideae were estimated to have diverged from each other during the Paleogene, in the Eocene era (ca. 40.63 Ma, HDP 35.45–45.16). Crown diversification of Cranichideae commenced in the late Eocene, ca. 35.8 Ma (HDP 29.54–41.28), with the divergence of subtribe Chloraeinae. Divergence between Pterostylidinae *s.s.* and the remainder of the tribe occurred in the early Oligocene, ca. 32.27 Ma (HDP 26.3–38.12) (Figure 5; Supplementary Material S3.2). Crown diversification of Pterostylidinae *s.s.* started in the Neogene period, during the mid-Miocene, ca. 14.7 Ma (HDP 10.59–19.27) with the divergence of clade A from the remainder of the genus. Divergence between clade B and clade C was also dated to the mid Miocene, ca. 12.98 Ma (HDP 9.25–46.88). Crown diversification of *Pterostylis* clades A, B, and C began in the late Miocene, with ca. 7.47 Ma (HDP 4.31–11.16) for clade A, 5.88 Ma (HDP 3.77–8.5) for clade B, and 10.43 (HDP 7.49–13.48) for clade C. All *Pterostylis* sections were estimated to have originated during the Miocene (Figure 4) and 92% of sampled *Pterostylis* species were estimated to have originated during the Quaternary (Figure 5; Supplementary Material S3.3).

**Figure 5.**
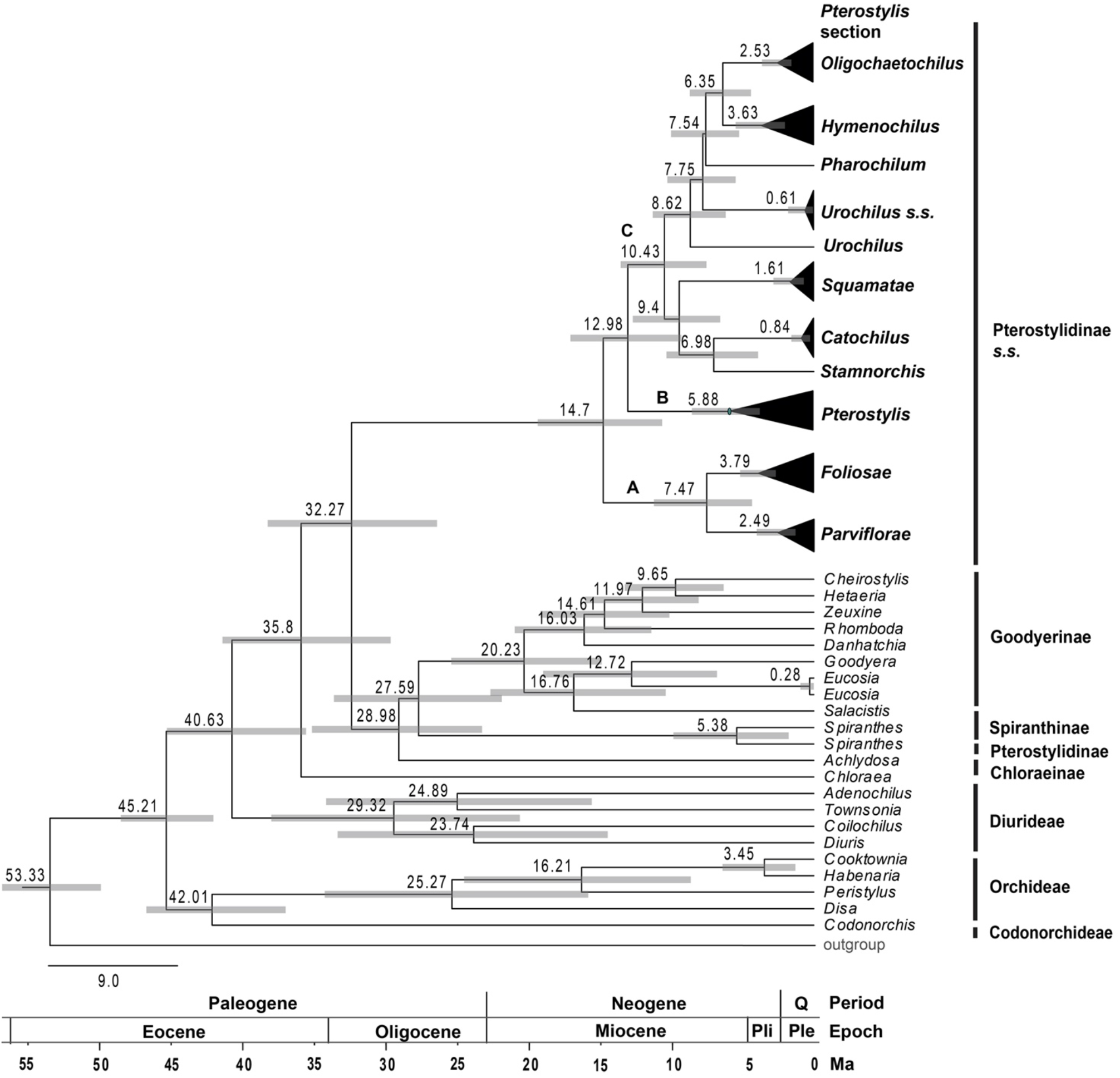
Chronogram showing divergence times of main lineages within Pterostylidinae *s.s.* and of tribes in Orchidoideae. Maximum clade credibility tree from Bayesian divergence time estimation based on 25 most informative plastid genes and an uncorrelated molecular clock model under the birth-death tree prior. Divergence times (Ma) are given at each node together with 95% highest posterior density (HDP) values indicated by grey bars. The full chronogram is provided in Supplementary Material S3.

#### Divergence time estimations based on nuclear dataset

The divergence time estimates based on the nuclear data consistently yielded older mean age estimates within Pterostylidinae *s.s.* than divergence dating based on the plastid data (Supplementary Material S3.4). The HDP intervals from the divergence time estimation based on the nuclear dataset were consistently wider than those from the inference based on the plastid data, thus indicating a larger uncertainty of the divergence estimates based on the nuclear data. The lower bounds of the HDP intervals for the age estimates derived from the nuclear data often approached the upper bounds of the HDP intervals of the plastid divergence dating analysis, however the means between the nuclear and plastid divergence ages lay consistently apart. For Pterostylidinae *s.s.* the divergence time estimates based on the nuclear dataset yielded a stem age of ca. 34.65 Ma (28.43–40.09) and a crown age of ca. 29.93 Ma (HDP 22.94–37.1). Crown ages for clades A and B were estimated to the early Miocene, ca. 22.69 Ma (HDP 14.96-30.90) and ca. 20.88 Ma (HDP 12.22-29.32), and to the late Oligocene with ca. 23.74 (1.44-32.06) for clade C. A comparison of divergence time estimates derived from the nuclear and the plastid dataset is provided in Supplementary Material S3.5.

### 3.3 Ancestral range estimation

Model testing of the three biogeographic models (DEC, DIVALIKE, BAYAREALIKE) using the Akaike information criterion identified the dispersal-extinction cladogenesis (DEC) model as best fit model for the ML estimation of ancestral ranges based on the chronogram derived from the plastid dataset (Supplementary Material 4.1).

Australia was inferred as the ancestral range of the most recent common ancestor (MRCA) of Pterostylidinae *s.s.*. As ancestral area of the subtribe, the Euronotian region received the highest relative probability (RP 0.72) (Figure 6). The Euronotian region was inferred as ancestral area of the MRCA of each of the three main clades, A (RP 0.72), B (RP 0.75), and C (RP 0.75). For the MRCAs of the following seven *Pterostylis* sections, the inferred ancestral area was also the Euronotian region: sect. *Foliosae* (RP 0.84), sect. *Hymenochilus* (RP 0.88), sect. *Oligochaetochilus* (RP 0.88), sect. *Parviflorae* (RP 0.84), sect. *Pharochilum* (RP 0.92), sect. *Pterostylis* (RP 0.75), and sect. *Squamatae* (RP 0.72) (Figure 6). A broader ancestral range comprised of the Euronotian region and southwest Australia was inferred for the MRCAs of sect. *Catochilus* (RP 0.87), sect. *Stamnorchis* (RP 0.87), sect. *Urochilus s.s.* (RP 0.63), and for the MRCA of *P. sargentii* (RP 0.89).

**Figure 6.**
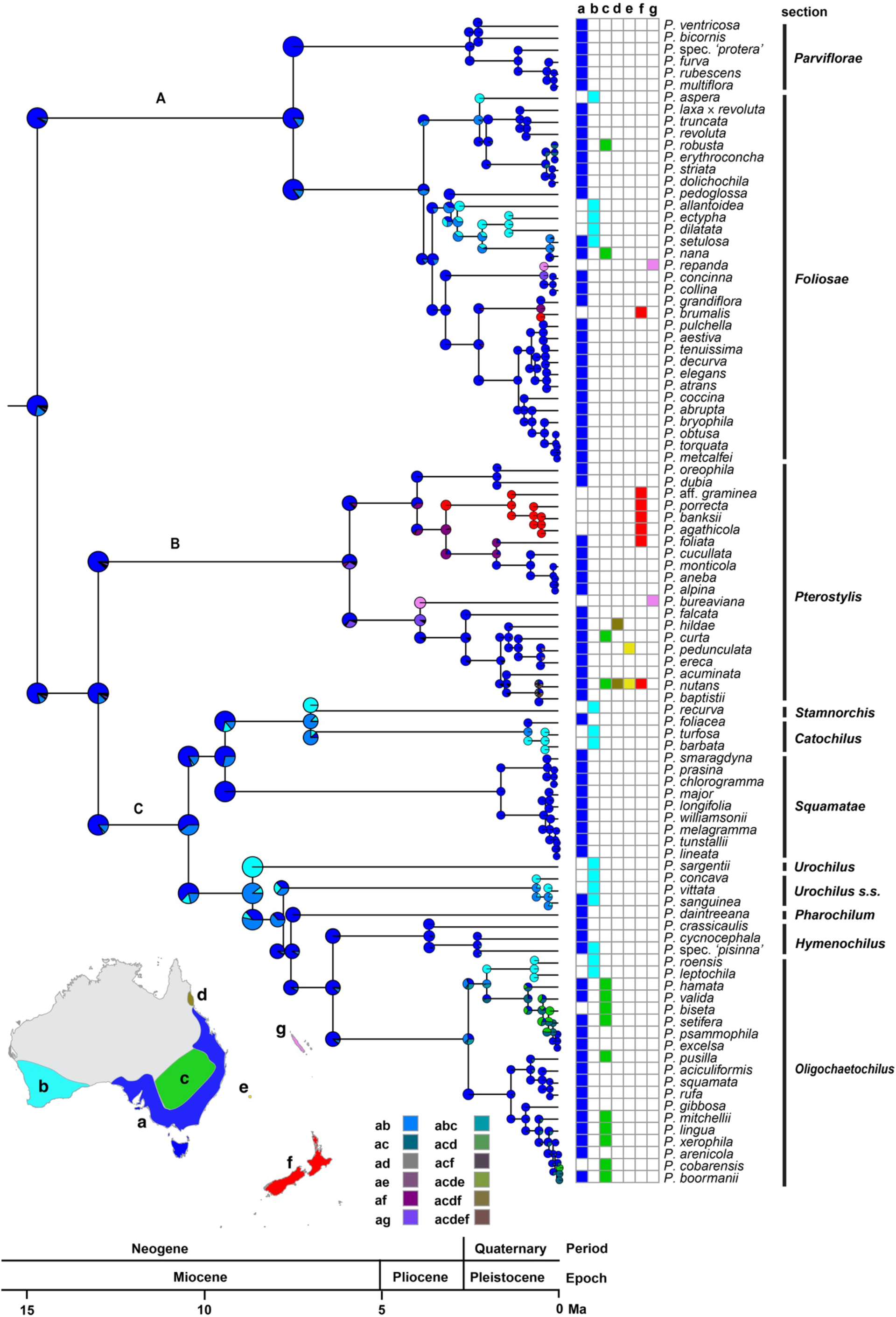
Ancestral range estimation in Pterostylidinae *s.s.*. Maximum likelihood estimation was carried out under the dispersal-extinction cladogenesis (DEC) model and used the maximum clade credibility tree from divergence dating analysis based on 25 plastid genes and an uncorrelated molecular clock model under the birth-death tree prior. Pie diagrams depict the relative probabilities of ancestral ranges. Relative probabilities of all alternative range scenarios are provided in Supplementary Material S4.2. Map insert shows area delineation and grid depicts area coding for each species. a: Euronotian region; b: southwest Australia; c: Eremaean region; d: Northern region (sub-region Atherton), e: Lord Howe Island; f: New Zealand; g: New Caledonia. The three major clades in *Pterostylis* are labelled above branches (as A, B, and C).

Relative probabilities of alternative range evolutionary scenarios are provided in Supplementary Material S.4.2.

Several independent range expansions and subsequent range shifts from the Euronotian region to southwest Australia were inferred. The earliest of these were inferred in clade C from the late Miocene onwards: in the MRCA of the clade comprising sections *Urochilus s.s., Pharochilum, Hymenochilus, Oligochaetochilus*, and *P. sargentii*, from ca. 8.6 Ma and in the MRCA of sections *Catochilus* and *Stamnorchis* from ca. 7.0 Ma onwards. Other range expansions and subsequent shifts from the Euronotian to southwest Australia were inferred to have occurred from the Pliocene and Pleistocene onwards: in sect. *Foliosae* (from ca. 3.0 Ma and 2.2 Ma), in sect. *Oligochaetochilus* (from ca. 2.5 Ma), and in sect. *Hymenochilus* (from ca. 2.3 Ma) (Figure 6).

At least ten range expansions from the Euronotian region to the adjacent Eremaean region were inferred, estimated to have occurred from the early Pleistocene onwards (Figure 6). At least two such range expansions were inferred in sect. *Foliosae* (from ca. 0.2 Ma and from 0.1 Ma) and in sect. *Pterostylis* (from ca. 1.1 Ma and ca. 0.5 Ma). At least six range expansions from the Euronotian region to the Eremaean region were inferred in sect. *Oligochaetochilus* commencing in the early to late Pleistocene, with at least two subsequent range shifts.

At least seven range expansions from continental Australia to other Australasian regions were inferred, in at least four instances with subsequent range shifts. Range expansions from continental Australia to New Zealand with subsequent range shifts were inferred to have occurred from the Euronotian region from the late Pliocene onwards in at least three instances, in sect. *Pterostylis* (from ca. 3.2 Ma) and in sect. *Foliosae* (from ca. 0.5 Ma). At least one range expansion to New Zealand (in sect. *Pterostylis*) was followed by *in situ* diversification. Range expansions from the Australian east coast to New Zealand and Lord Howe Island was inferred in at least one instance and occurred from the late Pleistocene onwards (in sect. *Pterostylis*). At least one range expansions from eastern Australia to Lorde Howe Island was inferred, estimated to have occurred from the late Pleistocene (ca. 0.5 Ma) (in sect. *Pterostylis*).

At least two range expansions from continental Australia to New Caledonia were inferred from the Euronotian region. The earliest of these range expansions was estimated to have occurred from the early Pliocene onwards (ca. 3.9 Ma) in sect. *Pterostylis*. The second range expansion to New Caledonia was estimated to have occurred from the mid Pleistocene onwards in sect. *Foliosae* (ca. 0.4 Ma) (Figure 6). Both inferred range expansions to New Caledonia were followed by range shifts. Relative probabilities for all range evolutionary scenarios are provided in Supplementary Material S4.2.

## 4 Discussion

### 4.1 Range evolution in Pterostylidinae

The divergence time estimates of our phylogenomic study provided further support for an Eocene origin and onset of crown diversification of Cranichideae, with comparable age estimates to Givnish et al. (2015) and Serna-Sánchez et al. (2021). A family-wide ancestral range reconstruction inferred a Neotropical-Australasian ancestral range for the MCRA for Cranichideae (Givnish et al., 2016). During the Eocene, Antarctica was still vegetated and provided biogeographic connections between these two major phytogeographic areas (Givnish et al., 2016).

Our study inferred an Australian origin of Pterostylidinae in the early Oligocene, ca. 32 Ma, coinciding with the timing of the complete separation of Australia from Antarctica and the establishment of the Antarctic Circumpolar Current at the Eocene–Oligocene boundary, which led to drastic climatic changes worldwide including global cooling and glaciation of Antarctica (Wilford, and Brown, 1994; Martin, 2006; Quilty, 1994). A previous ancestral range estimation (Givnish et al., 2016) also placed the origin of Pterostylidinae in the Oligocene (ca. 38 Ma) and inferred an Australian-Pacific or Australian origin of the lineage. However, Givnish et al. (2016) included only one representative for *Pterostylis* and applied a biogeographic area coding for the overall distribution of the genus. Our phylogenomic study with a broad sampling across the diversity within Pterostylidinae provided a more refined ancestral range estimation with well-supported evidence for an eastern Australian origin of the subtribe, in the Euronotian region.

The transition between the late Eocene and the early Oligocene with its stark climatic changes at global level was accompanied by profound vegetational changes in Australia which altered from a mosaic of mesotherm to megatherm rainforests with some sclerophyllous taxa to a predominance of cool temperate microtherm rainforests (Martin, 2006). Extant Pterostylidinae are most diverse and abundant in temperate habitats of Australia’s mesic biome and are particularly diverse in the Euronotian region of eastern Australia. The latter was inferred as the ancestral range of the subtribe, as well as of all three major lineages, the majority of *Pterostylis* sections, and of the majority of extant species. This points to a high degree of niche conservatism within Pterostylidinae. We assume that the wetter and cooler conditions during the early Oligocene already suited the environmental niche requirements of early Pterostylidinae. However, the predominance of dense cool temperate rainforest vegetation may still have restricted the availability of suitable, more open vegetation.

Our study estimated the onset of crown diversification of Pterostylidinae to the mid-Miocene, with emergence of the three major lineages (A, B, and C) in the Euronotian region. During the late Miocene, crown diversification of the three major lineages commenced and by the end of the Miocene, all lineages as recognised at sectional level by Janes and Duretto (2010) had emerged, the majority of these in the Euronotian region. During the late Miocene, the climate in Australia became increasingly dry, leading to severe vegetational changes. Rainforests considerably contracted, sclerophyll vegetation expanded, and a well-defined dry season established which facilitated regular burning (Kershaw et al., 1994, Martin, 2006). As geophytes with underground tubers, Pterostylidinae were well equipped for the shift to a seasonal climate with a more pronounced dry season, which occurred in the mid and late Miocene. In southeast Australia and southwest Australia, the expanding wet sclerophyll forests likely provided suitable habitat for Pterostylidinae, fostering range expansions and lineage divergence within the group. A meta-analysis based on dated molecular phylogenies of other Australian plant lineages (in Fabaceae, Myrtaceae, Casuarinaceae, and Protaceae) found an increase in speciation rates for lineages characteristic of sclerophyll habitats during the Miocene, coinciding with increased aridification and seasonality on the Australian continent (Crisp et al., 2004).

Our study found the earliest range expansions and shifts within Pterostylidinae commenced from the late Miocene onwards, with at least nine independent arrivals leading to the establishment of Pterostylidinae in southwest Australia, the earliest commencing from the late Miocene onwards (in MRCA of sect. *Stamnorchis* and remainder of clade from ca. 8.6 Ma, and in the MRCAs of sections *Catochilus* and *Stamnorchis* from ca. 7.0 Ma). By this time, the Nullarbor Plain, a karst surface constituting a strong edaphic barrier for many mesic plant species between southwest and southeast Australia, had already formed (Li et al., 2004). This implies that Pterostylidinae reached southwest Australia via long-distance dispersal of the wind-dispersed dust-like seeds across the Nullarbor Plain. Likewise, subsequent range expansions from the Euronotian region to southwest Australia occurring from the Pliocene and Pleistocene onwards, such as those found in sect. *Foliosae* and sect. *Oligochaetochilus*, can also be regarded a result of long-distance dispersal.

Our study revealed a remarkably recent origin of today’s species diversity in Pterostylidinae, with the majority of extant species estimated to have arisen during the Quaternary. This period saw a continued overall global cooling trend and increased aridification in Australia, leading to further expansion of open wood- and grasslands and continued decrease of dense forest cover (Wagstaff et al., 2001; Martin, 2006), resulting in an overall increase of suitable habitats for Pterostylidinae throughout the mesic zone of Australia. Rapid climatic oscillations of the Pleistocene led to multiple cycles of expansions and contractions of open wood- and grasslands versus dense forest vegetation during the drier glacial and moister interglacial cycles (Byrne et al., 2011; Martin, 2006). These cycles would have led to repeated fragmentation and expansion of suitable habitats for Pterostylidinae. Therefore, the climatic oscillations of the Pleistocene may have accelerated speciation in Pterostylidinae due to repeated cycles of genetic isolation of previously contiguous populations.

In several instances, range expansions from the Euronotian region into the adjacent, more arid Eremaean region were inferred to have commenced from the early Pleistocene onwards, and were most pronounced in sect. *Oligochaetochilus*. In more arid regions, Pterostylidinae are mostly found on well-drained sites and often in association with rocks, such as rock outcrops or domes, which allow the plants to grow in rock crevices and other situations where water run-off is concentrated, or in sandy soils where surrounding vegetation provides some shelter (Jones and Clements, 2002a). To ascertain to what extent the distributions in the Eremaean region may constitute relict populations or more recent dispersals to these pockets of suitable habitat, population genomic studies are warranted. Our study showed that today’s distribution of Pterostylidinae in the Pacific region, including Lord Howe Island, New Zealand, and New Caledonia, are of more recent origin, mostly from the early and mid-Pleistocene onwards, and therefore stem from long-distance dispersal events from eastern Australia. Our study also provided evidence for *in situ* diversification after long-distance dispersal to New Zealand. A spatio-temporal study in the Australasian orchid genus *Thelymitra* also found that extant distributions in the Pacific region were the result of long-distance dispersals from the Australian continent in more recent geological times, predominantly from eastern Australia, sometimes followed by speciation events (Nauheimer et al., 2018). Our results also support findings from an orchid-wide biogeographic study which identified Australia as important source area for migration to adjacent geographic regions (Givnish et al., 2016).

Our study found that age estimates from the nuclear dataset arrived at older age estimates, which could be due to the molecular clock and speciation model favoured in model testing (i.e., uncorrelated relaxed molecular clock and Yule model). In datasets with strong rate heterogeneity among lineages, especially in the presence of long stems and short crowns, the choice of the clock and speciation models can lead to bias in age estimates (Crisp et al. 2014; Sarver et al. 2019). Molecular studies based on nuclear phylogenomic data, such as derived through target enrichment, are desirable to further ascertain phylogenetic relationships and the spatio-temporal evolution in Pterostylidinae.

### 4.2 Evolutionary relationships in Cranichideae and Pterostylidinae and systematic implications

Our phylogenomic analysis of 75 plastid genes resolved subtribal relationships within Orchidoideae with maximum support, providing further evidence for the sister group relationship between Cranichideae and Diurideae, thus confirming previous molecular studies (Kores et al., 2001; Freudenstein et al., 2004; Givnish et al., 2015; Li et al., 2019; Serna-Sánchez et al., 2021; Perez-Escobar et al., 2021). Further, our study corroborated a sister group relationship between Codonorchideae and Orchideae, and the two in turn sister group to Cranichideae and Diurideae.

While early molecular studies based on two to four markers retrieved the same intertribal relationships (Kores et al., 2001; Freudenstein et al., 2004; Gustafson et al., 2010; Chomicki et al., 2015), previous plastid phylogenomic studies (Givnish et al., 2015; Serna-Sánchez et al., 2021) retrieved conflicting topologies, with either Orchideae as first diverging lineage, followed by Codonorchideae, then Diurideae and Cranichideae (Givnish et al., 2015) or Codonorchideae sister to Orchideae and the two in turn sister to Diurideae and Cranichideae (Serna-Sánchez et al., 2021). These studies exhibited lower support values for the position of Codonorchideae. Our study now provides strong support for the latter topology, in line with early molecular studies (Kores et al., 2001; Freudenstein et al., 2004; Gustafson et al., 2010; Chomicki et al., 2015). However, our phylogenetic analysis based on the nuclear rRNA cistron was unable to reconstruct intertribal relationships in Orchidoideae with confidence. Likewise, previous studies based on nuclear markers (ITS or *xdh*) were unable to resolve higher-level relationships in Orchidoideae (Clements et al., 2002; Gorniak et al., 2010). Hence, further studies based on an increased number of nuclear markers, such as those derived through target sequence capture, are warranted to further ascertain intertribal relationships in Orchidoideae.

Our study provided further molecular evidence for the phylogenetic placement of Pterostylidinae within Orchideae and for assessing the monophyly of Pterostylidinae.

Within Cranichideae, our study retrieved Chloraeinae as the first diverging lineage, followed by Pterostylidinae *s.s.*. The next diverging lineage comprised the monotypic genus *Achlydosa* as sister group to the remainder of Cranichideae. Previous molecular studies yielded conflicting results for the phylogenetic relationships among these lineages. A molecular study based on two plastid markers (*mat*K and *trn*L-F) (Kores et al., 2001) resolved Chloraeinae as diverging first, followed by Pterostylidinae, then *Achlydosa* (as *Megastylis glandulosa* (Schltr.) Schltr.), whereas a study based on four markers (*mat*K, *trn*L-F, *rbcL*, and ITS) showed Chloraeinae diverging first, followed by Pterostylidinae in sister group position to *Achlydosa* (as *Megastylis glandulosa*), as sister clade to the remainder of Cranichideae (Salazar et al., 2009). However, in the latter study, the results from each single marker yielded incongruent results for these lineages and in the combined analysis, the sister group relationship between Pterostylidinae *s.s.* and *Achlydosa* was not well supported. Our phylogenomic study provides support for phylogenetic relationships within Cranichideae as retrieved by Kores et al. (2001). Our phylogenomic study therefore does not support the taxonomic concept of Pterostylidinae *sensu* Chase et al. (2015) which included the genus *Achlydosa*. Chase et al. (2015) acknowledged that recognition of subtribe Achlydosinae may be warranted based on results of further studies. This study provides support for recognition of *Achlydosa* at subtribal level, as Achlydosinae *sensu* Clements et al. (2002).

Within Pterostylidinae *s.s.*, our study resolved relationships between the three major lineages in *Pterostylis* with strong support, with clade A as sister group to clades B and C. In previous phylogenetic studies based on ITS and/or *mat*K, relationships between the three major lineages remained unclear due to lack of resolution (Jones and Clements, 2002a) or low statistical support (Janes et al., 2010; Clements et al., 2011), showing either clade A as sister group to B and C or clades A and B as sister group to clade C. Based on the latter topology, Janes and Duretto (2010) proposed a subgeneric classification with two subgenera, *Pterostylis* (clades A and B) and *Oligochaetochilus* (clade C). Morphologically, the two subgenera were mainly differentiated by the position of the lateral sepals (deflexed in subgen. *Oligochaetochilus* with one exception (*P. recurva*) and erect in subgen. *Pterostylis*) and the presence/absence of barrier trichomes on the column wings (present in subgen. *Oligochaetochilus*, absent in subgen. *Pterostylis*). However, our study did not provide support for the monophyly of subgen. *Pterostylis sensu* Janes and Duretto (2010). A revision of the intrageneric classification of *Pterostylis s.l.* would require recognition of clade A at subgeneric level. Our study provided support for the monophyly of nine of the ten *Pterostylis* sections *sensu* Janes and Duretto (2010). However, the monophyly of sect. *Urochilus sensu* Janes and Duretto (2010) warrants further study as our results of the placement of *P. sargentii* remained ambiguous. In their circumscription of sect. *Urochilus*, Janes and Duretto (2010) broadened the original taxonomic concept of *Urochilus* to include *Ranorchis* (Jones and Clements, 2002b) due to lack of phylogenetic resolution in previous molecular studies. Should future studies support *P. sargentii* as a distinct lineage, the taxonomic classification of Jane and Duretto’s (2010) could be adjusted by adopting the original circumscriptions for *Urochilus* and *Ranorchis* for a revised sectional classification.

Based on the morphological distinctness of the lineages within Pterostylidinae, Jones and Clements (2002b) advocated for recognition of these groups at generic level. As illustrated in Figure 2, the sectional classification by Janes and Duretto (2010) and revised classification by Jones and Clements (Jones, 2015; Clements and Jones, 2016; Jones and Clements 2017; Jones, 2021) recognise the same morphological groups and evolutionary lineages (with the exception of sect. *Ranorchis* / *P.sargentii*), only at different taxonomic rank, and therefore are both equally well supported by our study. Further molecular study is required to clarify the taxonomic placement of *P. sargentii* and *P. daintreeana* due to remaining uncertainties regarding their phylogenetic position.

This study provided a phylogenomic framework for reassessing taxonomic concepts in the subtribe. The decision-making process to arrive at a taxonomic consensus in Pterostylidinae is complex as the key endeavour of systematics, to provide a useful natural classification, can be achieved in different ways. The endeavour to reflect new scientific insights in revised taxonomic classifications needs to be carefully weighed up against our aspiration to maintain taxonomic stability to provide a reliable and useful system to navigate biological diversity.

## Conclusions

This phylogenomic study clarified evolutionary relationships in Pterostylidinae and is the first to infer range evolution within the subtribe. The study provided well-supported evidence for an Australian origin of Pterostylidinae in the early Oligocene, after Australia fully separated from Antarctica. All main lineages in Pterostylidinae were inferred to have emerged during the Miocene when the Australian continent travelled to today’s geographic position and the continent underwent drastic vegetational changes in conjunction with increased aridification. This study showed that today’s species diversity is relatively young and largely originated during the Quaternary. Vegetational changes in conjunction with the climatic oscillations of the Pleistocene are seen as important drivers for the increase in diversification during the Quaternary. The Euronotian region, located in the eastern part of Australia’s mesic biome, was identified as ancestral area of the subtribe and as the area where Pterostylidinae predominantly underwent lineage diversification. The Euronotian region was further identified as key source area for other Australasian regions in the Pacific. Over its evolutionary history, Pterostylidinae remained largely confined to the mesic biome and hence exhibit a considerable degree of niche conservatism. This study provided an important phylogenomic framework for future studies on trait evolution in orchids, such as those based on morphological, anatomical or ecological traits, including pollination syndromes.

## 5 Data Availability Statement

The original contributions presented in the study are publicly available. This data can be found here: [https://www.ebi.ac.uk/ena/browser/text-search?query=PREJEB5249].

## 6 Author Contributions

KN, MC, and BM conceived and designed the study. LS and AM carried out laboratory work. KH, AM, SB, KN, LN, and LS performed analyses. KN, KH, MC, HZ, AM, LN, LS, and SB wrote the manuscript.

## 9 Acknowledgments

The authors wish to acknowledge the use of the Next Generation Sequencing services and facilities of the Australian Genomic Research Facilities (AGRF). Plant material was collected under kind permission of the Australian Capital Territory Government, PL201543 and PL201573; Government of South Australia’s Department of Environment, Water and Natural Resources, Y26092-1; the New South Wales National Parks & Wildlife Service, Scientific Licence, SL 100750; New South Wales State Forests, XX51121, HC 53443; Tasmanian Department of Primary Industry, Water and Environment, FL03121; the Victorian Government’s Department of Environment, Land, Water and Planning, Flora and Fauna Guarantee Act 1988, and National Parks Act 1975, 10006307; Government of Western Australian Department of Environment and Conservation, CE000418, NE002573, CE000160, SW015074; the Queensland Government’s Queensland Parks and Wildlife Service FO/000878/96/SAA, F1/000056/97/SAA, FO/000889/96/SAA, F1/00232/99/SAA, 1685, F1/000172/01/SAA and Queensland Government’s Department of Environment and Heritage Protection WITK11258712 and WISP11258812; and Nouvelle-Caledonie, Service de L’Environnement et de la Gestion des Parcs et Reserves (Authorisation no. 6024 765/ENV). We acknowledge G. Bradburn, P. Branwhite, C.P. Brock, W.H. Cherry, R. Collins, R. Crane, B. Dalyell, R. Datodi, P.J. de Lange, M. Duncan, A.R. Field, C.J. French, R.O. Gardner, B. Gray, W.K. Harris, T.N. Hayashi, D. Herd, D.L. Jones, R. Mawson, J. Moye, D.E. Murfet, W. Parr, E. Pisano, N. Pullman, N. Reiter, H.M.E. Richards, L.R. Roberts, J.W.D. Sawyer, A. Stephenson, R. Stockard, H. Wapstra, P.H. Weston, J. Whitfield, P. White, and M. Young for collection of plant material.

**Supplementary Material S1** | Plant material studied.

**Supplementary Material S2** | Phylogenetic relationships in Pterostylidinae based on plastid and nuclear data.

**Supplementary Material S3** | Divergence time estimations based on plastid and nuclear data. **Supplementary Material S4** | Model comparison of three biogeographic models for ancestral range estimations and relative probabilities of range evolutionary scenarios in Pterostylidinae.

