## Supplementary Material S1 for "Evolutionary relationships and range evolution of greenhood orchids (subtribe Pterostylidinae): insights from plastid phylogenomics"

Plant material studied. [1]Jones and Clements (2002), with revisions from Jones (2015), Clements & Jones (2016), and Jones & Clements (2017). *GenBank accession numbers of previously published DNA sequences. Herbarium acronyms: CANB, Australian National Herbarium; CHR: Allan Herbarium; CNS: Australian Tropical Herbarium; NSW: National Herbarium of New South Wales. Biogeographic area coding: a: Euronotian region; b: southwest Australia; c: Eremaean region; d: Northern region (sub-region Atherton), e: Lord Howe Island; f: New Zealand; g: New Caledonia. ENA: European Nucleotide Archive.

| Species | *Pterostylis* sections *sensu* Janes & Duretto (2010) | Species *sensu* Jones & Clements[1] | Voucher details | DNA extraction no. | ENA sample accession no. | Reference plastid data | Reference nuclear data | Area coding (abcdefg) |
|---|---|---|---|---|---|---|---|---|
| **Pterostylidinae** | | | | | | | | |
| *Achlydosa glandulosa* (Schltr.) M.A.Clem. & D.L.Jones | n/a | *Achlydosa glandulosa* (Schltr.) M.A.Clem. & D.L.Jones | M.A.Clements 11237 (CANB 950931.1) | CNS_G03414 | ERS11983043 | ERR9735413 | ERR9735413 | – |
| *Pterostylis abrupta* D.L.Jones | *Foliosae* G.Don | *Diplodium abruptum* (D.L.Jones) D.L.Jones & M.A.Clem | T.N.Hayashi 65 (CANB) | CNS_G04979 | ERS11983062 | ERR9735432 | ERR9735432 | 1000000 |
| *Pterostylis aciculiformis* (Nicholls) M.A.Clem. & D.L.Jones | *Oligochaetochilus* (Szlach.) Janes & Duretto | *Oligochaetochilus aciculiformis* (Nicholls) Szlach. | G.Bradburn 40 (CANB 892536.1) | CNS_G05943 | ERS11983063 | ERR9735433 | ERR9735433 | 1000000 |
| *Pterostylis acuminata* R.Br. | *Pterostylis* R.Br. | *Pterostylis acuminata* R.Br. | M.A.Clements 12014 (CANB 909845.1) | CNS_G03411 | ERS11983064 | ERR9735434 | – | 1000000 |
| *Pterostylis aestiva* D.L.Jones | *Foliosae* G.Don | *Diplodium aestivum* (D.L.Jones) D.L.Jones & M.A.Clem. | T.N.Hayashi 45 (CANB) | CNS_G05019 | ERS11983065 | ERR9735435 | ERR9735435 | 1000000 |
| *Pterostylis* aff. *graminea* "Kauri" | *Pterostylis* R.Br. | *Pterostylis* aff. *graminea* "Kauri" | B.P.J.Molloy 124/99 (CHR 531784) | CNS_G03985 | ERS11983066 | ERR9735436 | – | 0000010 |
| *Pterostylis agathicola* D.L.Jones, Molloy & M.A.Clem. | *Pterostylis* R.Br. | *Pterostylis agathicola* D.L.Jones, Molloy & M.A.Clem. | P.J.deLange 3903 & R.O.Gardner 082/99 (CHR 518307) | CNS_G06188 | ERS11983067 | ERR9735437 | – | 0000010 |
| *Pterostylis allantoidea* R.S.Rogers | *Foliosae* G.Don | *Diplodium allantoideum* (R.S.Rogers) D.L.Jones & M.A.Clem. | M.A.Clements 10891a (CANB 644580.1) | CNS_G06192 | ERS11983068 | ERR9735438 | ERR9735438 | 0100000 |
| *Pterostylis alpina* R.S.Rogers | *Pterostylis* R.Br. | *Pterostylis alpina* R.S.Rogers | ORG7428 (CANB 950938.1) | CNS_G05940 | ERS11983069 | ERR9735439 | ERR9735439 | 1000000 |
| *Pterostylis aneba* D.L.Jones | *Pterostylis* R.Br. | *Pterostylis aneba* R.S.Rogers | P. Branwhite 392 (CANB 950935.1) | CNS_G03408 | ERS11983070 | ERR9735440 | ERR9735440 | 1000000 |
| *Pterostylis arenicola* M.A.Clem. & J.Stewart | *Oligochaetochilus* (Szlach.) Janes & Duretto | *Oligochaetochilus arenicola* (M.A.Clem. & J.Stewart) Szlach. | M.Young 29 (CANB 950922.1) | CNS_G05951 | ERS11983071 | ERR9735441 | – | 1000000 |
| *Pterostylis aspera* D.L.Jones & M.A.Clem. | *Foliosae* G.Don | *Diplodium asperum* (D.L.Jones & M.A.Clem.) D.L.Jones & M.A.Clem. | C.J.French 6115 (CANB 666990.1) | CNS_G06194 | ERS11983072 | ERR9735442 | ERR9735442 | 0100000 |
| *Pterostylis atrans* D.L.Jones | *Foliosae* G.Don | *Diplodium atrans* (D.L.Jones) D.L.Jones & M.A.Clem. | T.N.Hayashi 38 (CANB) | CNS_G04407 | ERS11983073 | ERR9735443 | ERR9735443 | 1000000 |
| *Pterostylis atrosanguinea* (D.L.Jones & C.J.French) D.L.Jones & C.J.French | *Urochilus* (D.L.Jones & M.A.Clem.) Janes & Duretto | *Urochilus atrosanguineus* D.L.Jones & C.J.French | C.J.French 6120 (CANB 666994.1) | CNS_G06186 | ERS11983074 | – | ERR9735444 | – |
| *Pterostylis banksii* R.Br. ex A.Cunn. | *Pterostylis* R.Br. | *Pterostylis banksii* A.Cunn. | W.Parr 087/99 (CHR 518312) | CNS_G04072 | ERS11983075 | ERR9735445 | – | 0000010 |
| *Pterostylis baptistii* Fitzg. | *Pterostylis* R.Br. | *Pterostylis baptistii* Fitzg. | R.Crane 2088 (CANB 665689.1) | CNS_G04034 | ERS11983076 | ERR9735446 | – | 1000000 |
| *Pterostylis barbata* Lindl. | *Catochilus* Benth. | *Plumatichilos barbata* Szlach. | M.A.Clements 11966B (CANB 891201.1) | CNS_G01354 | ERS11983077 | ERR9735447 | this study | 0100000 |

| Species | *Pterostylis* sections *sensu* Janes & Duretto (2010) | Species *sensu* Jones & Clements[1] | Voucher details | DNA extraction no. | ENA sample accession number | Reference plastid data | Reference nuclear data | Area coding (abcdefg) |
|---|---|---|---|---|---|---|---|---|
| *Pterostylis bicornis* D.L.Jones & M.A.Clem. | *Parviflorae* (Bent.) Janes & Duretto | *Petrorchis bicornis* (D.L.Jones & M.A.Clem.) D.L.Jones & M.A.Clem. | B.Dalyell ORG 4865 (CANB 673100.1) | CNS_G06427 | ERS11983078 | ERR9735448 | ERR9735448 | 1000000 |
| *Pterostylis boormanii* Rupp | *Oligochaetochilus* (Szlach.) Janes & Duretto | *Oligochaetochilus boormanii* (Rupp) Szlach. | T.N.Hayashi 112 (CANB 950907.1) | CNS_G05972 | ERS11983079 | ERR9735449 | ERR9735449 | 1010000 |
| *Pterostylis brumalis* L.B.Moore | *Foliosae* G.Don | *Diplodium brumale* (L.B.Moore) D.L.Jones, Molloy & M.A.Clem. | N.Pullman 078/99 (CHR 518302) | CNS_G06189 | ERS11983080 | ERR9735450 | ERR9735450 | 0000010 |
| *Pterostylis bryophila* D.L.Jones | *Foliosae* G.Don | *Diplodium bryophilum* (D.L.Jones) D.L.Jones & M.A.Clem. | D.E.Murfet 1772 (CANB 676620.1) | CNS_G06413 | ERS11983081 | ERR9735451 | ERR9735451 | 1000000 |
| *Pterostylis bureaviana* Schltr. | *Pterostylis* R.Br. | *Pterostylis bureaviana* Schltr. | M.A.Clements 11225 (CANB 664441.1) | CNS_G06209 | ERS11983082 | ERR9735452 | – | 0000001 |
| *Pterostylis bureaviana* Schltr. | *Pterostylis* R.Br. | *Pterostylis bureaviana* Schltr. | M.A.Clements 11180 (CANB 664439.1) | CNS_G06191 | ERS11983083 | ERR9735453 | ERR9735453 | – |
| *Pterostylis chlorogramma* D.L.Jones & M.A.Clem. | *Squamatae* G.Don | *Bunochilus chlorogrammus* (D.L.Jones & M.A.Clem.) D.L.Jones & M.A.Clem. | T.N.Hayashi 97 (CANB 891065.1) | CNS_G05969 | ERS11983084 | ERR9735454 | ERR9735454 | 1000000 |
| *Pterostylis cobarensis* M.A.Clem. | *Oligochaetochilus* (Szlach.) Janes & Duretto | *Oligochaetochilus cobarensis* (M.A.Clem.) Szlach. | G.Bradburn 27B (CANB 950910.1) | CNS_G05946 | ERS11983085 | ERR9735455 | ERR9735455 | 0010000 |
| *Pterostylis coccina* Fitzg. | *Foliosae* G.Don | *Diplodium coccinum* (Fitzg.) D.L.Jones & M.A.Clem. | G.Bradburn 11 (CANB) | CNS_G04998 | ERS11983086 | ERR9735456 | ERR9735456 | 1000000 |
| *Pterostylis collina* (Rupp) M.A.Clem. & D.L.Jones | *Foliosae* G.Don | *Diplodium collinum* (Rupp) M.A.Clem. & D.L.Jones | M.A.Clements 12419 (CANB) | CNS_G06196 | ERS11983087 | ERR9735456 | ERR9735456 | 1000000 |
| *Pterostylis concava* D.L.Jones & M.A.Clem. | *Urochilus* (D.L.Jones & M.A.Clem.) Janes & Duretto | *Urochilus concavus* (D.L.Jones & M.A.Clem.) D.L.Jones & M.A.Clem. | C.J.French 6119 (CANB 666993.1) | CNS_G06212 | ERS11983088 | ERR9735456 | ERR9735456 | 0100000 |
| *Pterostylis concinna* R.Br. | *Foliosae* G.Don | *Diplodium concinnum* (R.Br.) M.A.Clem. & D.L.Jones | M.A.Clements 12408 (CANB) | CNS_G06197 | ERS11983089 | ERR9735459 | ERR9735459 | 1000000 |
| *Pterostylis crassicaulis* (D.L.Jones & M.A.Clem.) G.N.Backh. | *Hymenochilus* (D.L.Jones & M.A.Clem.) Janes & Duretto | *Hymenochilus crassicaulis* D.L.Jones & M.A.Clem. | M.A.Clements 12333 (CANB 950934.1) | CNS_G05968 | ERS11983090 | ERR9735460 | ERR9735460 | 1000000 |
| *Pterostylis cucullata* R.Br. | *Pterostylis* R.Br. | *Pterostylis cucullata* R.Br. | R.Mawson ORG5141 (CANB 677099.1) | CNS_G06211 | ERS11983091 | ERR9735461 | ERR9735461 | 1000000 |
| *Pterostylis curta* R.Br. | *Pterostylis* R.Br. | *Pterostylis curta* R.Br. | ORG7345 (CANB 950929.1) | CNS_G05071 | ERS11983091 | – | ERR9735461 | – |
| *Pterostylis curta* R.Br. | *Pterostylis* R.Br. | *Pterostylis curta* R.Br. | R.Crane 2042 (CANB 665695.1) | CNS_G03412 | ERS11983093 | ERR9735463 | ERR9735463 | 1010000 |
| *Pterostylis cycnocephala* Fitzg. | *Hymenochilus* (D.L.Jones & M.A.Clem.) Janes & Duretto | *Hymenochilus cymbellus* Jones ined. | M.A.Clements 11916 (CANB 906118.1) | CNS_G02745 | ERS11983094 | ERR9735464 | ERR9735464 | 1000000 |
| *Pterostylis daintreeana* F.Muell. ex Benth. | *Pharochilum* (D.L.Jones & M.A.Clem.) Janes & Duretto | *Pharochilum daintreeanum* (Benth.) D.L.Jones & M.A.Clem | T.N.Hayashi 76 (CANB 950936.1) | CNS_G05043 | ERS11983095 | ERR9735465 | ERR9735465 | 1000000 |

| Species | *Pterostylis* sections *sensu* Janes & Duretto (2010) | Species *sensu* Jones & Clements[1] | Voucher details | DNA extraction no. | ENA sample accession number | Reference plastid data | Reference nuclear data | Area coding (abcdefg) |
|---|---|---|---|---|---|---|---|---|
| *Pterostylis decurva* R.S.Rogers | *Foliosae* G.Don | *Diplodium decurvum* (R.S.Rogers) D.L.Jones & M.A.Clem. | M.A.Clements 10744 (CANB 664394.1) | CNS_G05977 | ERS11983096 | ERR9735466 | ERR9735466 | 1000000 |
| *Pterostylis dilatata* A.S.George | *Foliosae* G.Don | *Diplodium dilatatum* (A.S.George) D.L.Jones & M.A.Clem. | C.J.French 6116 (CANB 666991.1) | CNS_G06193 | ERS11983097 | ERR9735467 | ERR9735467 | 0100000 |
| *Pterostylis dolichochila* D.L.Jones & M.A.Clem. | *Foliosae* G.Don | *Diplodium dolichochilum* (D.L.Jones & M.A.Clem.) D.L.Jones & M.A.Clem. | M.A.Clements 11922 (CANB 909820.1) | CNS_G05959 | ERS11983098 | ERR9735468 | ERR9735468 | 1000000 |
| *Pterostylis dubia* R.Br. | *Pterostylis* R.Br. | *Pterostylis dubia* R.Br. | M.A.Clements 11158 (CANB 681498.1) | CNS_G06185 | ERS11983099 | ERR9735469 | ERR9735469 | 1000000 |
| *Pterostylis ectypha* (D.L.Jones & C.J.French) D.L.Jones & C.J.French | *Foliosae* G.Don | *Diplodium ectyphum* D.L.Jones & C.J.French | C.J.French 1570 (CANB 624821.1) | CNS_G02787 | ERS11983100 | ERR9735470 | ERR9735470 | 0100000 |
| *Pterostylis elegans* D.L.Jones | *Foliosae* G.Don | *Diplodium elegans* (D.L.Jones) D.L.Jones & M.A.Clem. | T.N.Hayashi 72 (CANB) | CNS_G05029 | ERS11983101 | ERR9735471 | ERR9735471 | 1000000 |
| *Pterostylis erecta* T.E.Hunt | *Pterostylis* R.Br. | *Pterostylis erecta* T.E.Hunt | M.A.Clements 11512 (CANB 882081.1) | CNS_G06198 | ERS11983102 | ERR9735472 | ERR9735472 | 1000000 |
| *Pterostylis erythroconcha* M.A.Clem. & D.L.Jones | *Foliosae* G.Don | *Diplodium erythroconchum* (M.A.Clem. & D.L.Jones) D.L.Jones & M.A.Clem. | M.Young 17 (CANB 950923.1) | CNS_G05942 | ERS11983103 | ERR9735473 | ERR9735473 | 1000000 |
| *Pterostylis excelsa* M.A.Clem. | *Oligochaetochilus* (Szlach.) Janes & Duretto | *Oligochaetochilus excelsus* (M.A.Clem.) Szlach. | M.A.Clements 11889 (CANB 950919.1) | CNS_G02748 | ERS11983104 | ERR9735474 | ERR9735474 | 1000000 |
| *Pterostylis falcata* R.S.Rogers | *Pterostylis* R.Br. | *Pterostylis falcata* R.S.Rogers | T.N.Hayashi 25 (CANB 950906.1) | CNS_G04423 | ERS11983105 | ERR9735475 | ERR9735475 | 1000000 |
| *Pterostylis foliacea* (D.L.Jones) D.L.Jones | *Catochilus* Benth. | *Plumatichilos foliaceus* D.L.Jones | M.A.Clements 11896 (CANB 906103.1) | CNS_G02746 | ERS11983106 | ERR9735476 | ERR9735476 | 1000000 |
| *Pterostylis foliata* Hook.f. | *Pterostylis* R.Br. | *Pterostylis gracilis* Nicholls | M.A.Clements 11797 (CANB 906077.1) | CNS_G06207 | ERS11983107 | ERR9735477 | ERR9735477 | 1000010 |
| *Pterostylis furva* (D.L.Jones) D.L.Jones | *Parviflorae* (Bent.) Janes & Duretto | *Speculantha furva* D.L.Jones | T.N.Hayashi 54 (CANB 950913.1) | CNS_G05034 | ERS11983108 | ERR9735478 | ERR9735478 | 1000000 |
| *Pterostylis gibbosa* R.Br. | *Oligochaetochilus* (Szlach.) Janes & Duretto | *Oligochaetochilus gibbosus* (R.Br.) Szlach. | M.A.Clements 11039 (CANB 655142.1) | CNS_G01273 | ERS11983109 | ERR9735479 | ERR9735479 | 1000000 |
| *Pterostylis grandiflora* R.Br. | *Foliosae* G.Don | *Diplodium grandiflorum* (R.Br.) D.L.Jones & M.A.Clem | M.A.Clements 12010 (CANB 909841.1) | CNS_G04878 | ERS11983110 | ERR9735480 | – | 1000000 |
| *Pterostylis hamata* Blackmore & Clemesha | *Oligochaetochilus* (Szlach.) Janes & Duretto | *Oligochaetochils hamatus* (Blackmore & Clemesha) Szlach. | G.Bradburn 25 (CANB 950909.1) | CNS_G05980 | ERS11983111 | ERR9735481 | ERR9735481 | 1010000 |
| *Pterostylis hildae* Nicholls | *Pterostylis* R.Br. | *Pterostylis hildae* Nicholls | B.Dalyell ORG3471 (CANB 629210.1) | CNS_G04114 | ERS11983112 | ERR9735482 | ERR9735482 | 1001000 |
| *Pterostylis laxa* Blackmore x *revoluta* R.Br. | *Foliosae* G.Don | *Diplodium laxum* (Blackmore) D.L.Jones & M.A.Clem. x *revolutum* (R.Br.) D.L.Jones & M.A.Clem. | M.A.Clements 12093 (CANB) | CNS_G05996 | ERS11983113 | ERR9735483 | ERR9735483 | 1000000 |
| *Pterostylis leptochila* M.A.Clem. & D.L.Jones | *Oligochaetochilus* (Szlach.) Janes & Duretto | *Oligochateochilus leptochilus* (M.A.Clem. & D.L.Jones) Szlach. | M.A.Clements 12235 & L. Nauheimer (CANB) | CNS_G05964 | ERS11983114 | ERR9735484 | ERR9735484 | 0100000 |
| *Pterostylis lineata* (D.L.Jones) G.N.Backh. | *Squamatae* G.Don | *Bunochilus lineatus* D.L.Jones | D.L.Jones 16514 (CANB 607105.1) | CNS_G05978 | ERS11983115 | ERR9735485 | ERR9735485 | 1000000 |

| **Species** | ***Pterostylis* sections *sensu* Janes & Duretto (2010)** | **Species *sensu* Jones & Clements[1]** | **Voucher details** | **DNA extraction no.** | **ENA sample accession number** | **Reference plastid data** | **Reference nuclear data** | **Area coding (abcdefg)** |
|---|---|---|---|---|---|---|---|---|
| *Pterostylis lingua* M.A.Clem. | *Oligochaetochilus* (Szlach.) Janes & Duretto | *Oligochaetochilus linguus* (M.A.Clem.) Szlach. | N.Reiter ORG6356 (CANB) | CNS_G06202 | ERS11983116 | ERR9735486 | ERR9735486 | 1010000 |
| *Pterostylis longifolia* R.Br. | *Squamatae* G.Don | *Bunochilus longifolius* (R.Br.) D.L.Jones & M.A.Clem. | M.A.Clements 11755 (CANB 751196.1) | CNS_G04968 | ERS11983117 | ERR9735487 | ERR9735487 | 1000000 |
| *Pterostylis major* (D.L.Jones) G.N.Backh. | *Squamatae* G.Don | *Bunochilus major* D.L.Jones | T.N.Hayashi 83 (CANB 891072.1) | CNS_G05947 | ERS11983118 | ERR9735488 | ERR9735488 | 1000000 |
| *Pterostylis melagramma* D.L.Jones | *Squamatae* G.Don | *Bunochilus melagrammus* (D.L.Jones) D.L.Jones & M.A.Clem. | M.A.Clements 10738 (CANB 664392.1) | CNS_G04077 | ERS11983119 | ERR9735489 | ERR9735489 | 1000000 |
| *Pterostylis metcalfei* D.L.Jones | *Foliosae* G.Don | *Diplodium metcalfei* (D.L.Jones) D.L.Jones & M.A.Clem. | T.N.Hayashi 82 (CANB) | CNS_G05979 | ERS11983120 | ERR9735490 | ERR9735490 | 1000000 |
| *Pterostylis mitchellii* Lindl. | *Oligochaetochilus* (Szlach.) Janes & Duretto | *Oligochaetochilus mitchellii* (Lindl.) Szlach. | G.Bradburn 24 (CANB 950916.1) | CNS_G05953 | ERS11983121 | ERR9735491 | ERR9735491 | 1010000 |
| *Pterostylis monticola* D.L.Jones | *Pterostylis* R.Br. | *Pterostylis monticola* D.L.Jones | T.N.Hayashi 18 (CANB 950904.1) | CNS_G04391 | ERS11983122 | ERR9735492 | ERR9735492 | 1000000 |
| *Pterostylis multiflora* (D.L.Jones) G.N.Backh. | *Parviflorae* (Bent.) Janes & Duretto | *Speculantha multiflora* D.L.Jones | T.N.Hayashi 47 (CANB 950905.1) | CNS_G05016 | ERS11983123 | ERR9735493 | ERR9735493 | 1000000 |
| *Pterostylis nana* R.Br. | *Foliosae* G.Don | *Diplodium clavigerum* (Fitzg.) D.L.Jones & M.A.Clem. | G.Bradburn 49 (CANB 950915.1) | CNS_G06206 | ERS11983124 | ERR9735494 | ERR9735494 | 1010000 |
| *Pterostylis nutans* R.Br. | *Pterostylis* R.Br. | *Pterostylis nutans* R.Br. | M.A.Clements 12119 (CANB 906133.1) | CNS_G04366 | ERS11983126 | ERR9735496 | ERR9735496 | 1011010 |
| *Pterostylis nutans* R.Br. | *Pterostylis* R.Br. | *Pterostylis hispidula* Fitzg. | K.Schulte 116 (CANB 950908.1) | CNS_G00238 | ERS11983125 | ERR9735495 | ERR9735495 | – |
| *Pterostylis obtusa* R.Br. | *Foliosae* G.Don | *Diplodium obtusum* (R.Br.) D.L.Jones & M.A.Clem. | G.Bradburn 14 (CANB) | CNS_G05024 | ERS11983127 | ERR9735497 | ERR9735497 | 1000000 |
| *Pterostylis oreophila* Clemesha | *Pterostylis* R.Br. | *Pterostylis oreophila* Clemesha | M.A.Clements 12335 (CANB) | CNS_G05966 | ERS11983128 | ERR9735498 | ERR9735498 | 1000000 |
| *Pterostylis pedoglossa* Fitzg. | *Foliosae* G.Don | *Diplodium pedoglossum* (Fitzg.) M.A.Clem. & D.L.Jones | T.N.Hayashi 78 (CANB) | CNS_G05044 | ERS11983129 | ERR9735499 | ERR9735499 | 1000000 |
| *Pterostylis pedunculata* R.Br. | *Pterostylis* R.Br. | *Pterostylis pedunculata* R.Br. | M.A.Clements 12116 (CANB 950914.1) | CNS_G04387 | ERS11983130 | ERR9735500 | – | 1000100 |
| *Pterostylis porrecta* D.L.Jones, Molloy & M.A.Clem. | *Pterostylis* R.Br. | *Pterostylis porrecta* D.L.Jones, Molloy & M.A.Clem. | P.J.deLange & J.W.D.Sawyer 149/99 (CHR 531809) | CNS_G06190 | ERS11983131 | ERR9735501 | – | 0000010 |
| *Pterostylis prasina* (D.L.Jones) G.N.Backh. | *Squamatae* G.Don | *Bunochilus prasinus* D.L.Jones | T.N.Hayashi 104 (CANB 950927.1) | CNS_G05933 | ERS11983132 | ERR9735502 | ERR9735502 | 1000000 |
| *Pterostylis psammophilus* (D.L.Jones) R.J.Bates | *Oligochaetochilus* (Szlach.) Janes & Duretto | *Oligochaetochilus psammophilus* D.L.Jones | M.Young 22 (CANB 950921.1) | CNS_G05967 | ERS11983133 | ERR9735503 | ERR9735503 | 1000000 |
| *Pterostylis pulchella* Messmer | *Foliosae* G.Don | *Diplodium pulchellum* (Messmer) D.L.Jones & M.A.Clem. | H.M.E. Richards & R. Datodi 617 (CANB 950933.1) | CNS_G04872 | ERS11983134 | ERR9735504 | ERR9735504 | 1000000 |
| *Pterostylis pusilla* R.S.Rogers | *Oligochaetochilus* (Szlach.) Janes & Duretto | *Oligochaetochilus pusillus* (R.S.Rogers) Szlach. | M.A.Clements 11872 (CANB 950920.1) | CNS_G02747 | ERS11983135 | ERR9735505 | – | 1010000 |
| *Pterostylis pyramidalis* Lindl. | *Foliosae* G.Don | *Diplodium pyramidialis* Lindl.) D.L.Jones & M.A.Clem. | M.A.Clements 11940B (CANB 891175.1) | CNS_G01115 | ERS11983136 | – | ERR9735506 | – |

| Species | *Pterostylis* sections *sensu* Janes & Duretto (2010) | Species *sensu* Jones & Clements[1] | Voucher details | DNA extraction no. | ENA sample accession number | Reference plastid data | Reference nuclear data | Area coding (abcdefg) |
|---|---|---|---|---|---|---|---|---|
| *Pterostylis recurva* Benth. | *Stamnorchis* (D.L.Jones & M.A.Clem.) Janes & Duretto | *Stamnorchis recurva* (Benth.) D.L.Jones & M.A.Clem. | M.A.Clements 11938 (CANB 891173.1) | CNS_G01355 | ERS11983137 | ERR9735507 | ERR9735507 | 0100000 |
| *Pterostylis repanda* (M.A.Clem. & D.L.Jones) J.M.H.Shaw | *Foliosae* G.Don | *Diplodium repandum* M.A.Clem. & D.L.Jones | D.L.Jones 15577 (CBG 9908875.1) | CNS_G04883 | ERS11983138 | ERR9735508 | ERR9735508 | 0000001 |
| *Pterostylis repanda* (M.A.Clem. & D.L.Jones) J.M.H.Shaw | *Foliosae* G.Don | *Diplodium repandum* M.A.Clem. & D.L.Jones | M.A.Clements 11213 (CANB 664440.1) | CNS_G06210 | ERS11983139 | ERR9735509 | ERR9735509 | – |
| *Pterostylis revoluta* R.Br. | *Foliosae* G.Don | *Diplodium revolutum* (R.Br.) D.L.Jones & M.A.Clem. | G.Bradburn 12 (CANB 950911.1) | CNS_G04997 | ERS11983140 | ERR9735510 | ERR9735510 | 1000000 |
| *Pterostylis robusta* R.S.Rogers | *Foliosae* G.Don | *Diplodium robustum* (R.S.Rogers) D.L.Jones & M.A.Clem. | M.A.Clements 11890 (CANB 906098.1) | CNS_G03422 | ERS11983141 | ERR9735511 | ERR9735511 | 1010000 |
| *Pterostylis roensis* M.A.Clem. & D.L.Jones | *Oligochaetochilus* (Szlach.) Janes & Duretto | *Oligochaetochilus roensis* (M.A.Clem. & D.L.Jones) Szlach. | M.A.Clements 10893b (CANB 644582.1) | CNS_G06204 | ERS11983142 | ERR9735512 | ERR9735512 | 0100000 |
| *Pterostylis rubescens* (D.L.Jones) G.N.Backh. | *Parviflorae* (Bent.) Janes & Duretto | *Speculantha rubescens* D.L.Jones | M.A.Clements 11847 (CANB 906086.1) | CNS_G05012 | ERS11983143 | ERR9735513 | ERR9735513 | 1000000 |
| *Pterostylis rufa* R.Br. | *Oligochaetochilus* (Szlach.) Janes & Duretto | *Oligochaetochilus rufus* (R.Br.) Szlach. | D.Herd ORG5019 (CANB 672882.1) | CNS_G06187 | ERS11983144 | ERR9735514 | ERR9735514 | 1000000 |
| *Pterostylis sanguinea* D.L.Jones & M.A.Clem. | *Urochilus* (D.L.Jones & M.A.Clem.) Janes & Duretto | *Urochilus sanguineus* (D.L.Jones & M.A.Clem.) D.L.Jones & M.A.Clem. | T.N.Hayashi 100 (CANB 950926.1) | CNS_G05945 | ERS11983145 | ERR9735515 | ERR9735515 | 1100000 |
| *Pterostylis sargentii* C.R.P.Andrews | *Urochilus* (D.L.Jones & M.A.Clem.) Janes & Duretto | *Ranorchis sargentii* (C.R.P.Andrews) D.L.Jones & M.A.Clem. | M.A.Clements 12308 & L.Nauheimer (CANB 950928.1) | CNS_G05973 | ERS11983146 | ERR9735516 | ERR9735516 | 0100000 |
| *Pterostylis setifera* M.A.Clem., Matthias & D.L.Jones | *Oligochaetochilus* (Szlach.) Janes & Duretto | *Oligochaetochilus setifer* (M.A.Clem., Matthias & D.L.Jones) Szlach. | G.Bradburn 27 (CANB 950937.1) | CNS_G05938 | ERS11983147 | ERR9735517 | ERR9735517 | 1010000 |
| *Pterostylis setulosa* (D.L.Jones & C.J.French) D.L.Jones & C.J.French | *Foliosae* G.Don | *Diplodium setulosum* D.L.Jones & C.J.French | M.A.Clements 10852 (CANB 644541.1) | CNS_G04435 | ERS11983148 | ERR9735518 | ERR9735518 | 1100000 |
| *Pterostylis smaragdyna* D.L.Jones & M.A.Clem. | *Squamatae* G.Don | *Bunochilus smaragdynus* (D.L.Jones & M.A.Clem.) D.L.Jones & M.A.Clem. | T.N.Hayashi 98 (CANB 891075.1) | CNS_G05958 | ERS11983149 | ERR9735519 | ERR9735519 | 1000000 |
| *Pterostylis* spec. *'pisinna'* | *Hymenochilus* (D.L.Jones & M.A.Clem.) Janes & Duretto | *Hymenochilus pisinnus* D.L.Jones ined. | M.Young 10a (CANB 950924.1) | CNS_G05970 | ERS11983150 | ERR9735520 | ERR9735520 | 1100000 |
| *Pterostylis* spec. *'pisinna'* | *Hymenochilus* (D.L.Jones & M.A.Clem.) Janes & Duretto | *Hymenochilus pisinnus* D.L.Jones ined. | M.A.Clements 11920 (CANB 909819.1) | CNS_G02754 | ERS11983151 | ERR9735521 | ERR9735521 | – |
| *Pterostylis* spec. *'protera'* | *Parviflorae* (Bent.) Janes & Duretto | *Speculantha* spec. *'protera'* | D.L.Jones ORG 7427 (CANB) | CNS_G05963 | ERS11983152 | ERR9735522 | – | 1000000 |
| *Pterostylis squamata* R.Br. | *Oligochaetochilus* (Szlach.) Janes & Duretto | *Oligochaetochilus squamatus* (R.Br.) Szlach. | T.N.Hayashi 30 (CANB) | CNS_G04429 | ERS11983153 | ERR9735523 | ERR9735523 | 1000000 |

| Species | *Pterostylis* sections *sensu* Janes & Duretto (2010) | Species *sensu* Jones & Clements[1] | Voucher details | DNA extraction no. | ENA sample accession number | Reference plastid data | Reference nuclear data | Area coding (abcdefg) |
|---|---|---|---|---|---|---|---|---|
| *Pterostylis striata* Fitzg. | *Foliosae* G.Don | *Diplodium striatum* (Fitzg.) D.L.Jones & M.A.Clem. | M.A. Clements 11782 (CANB) | CNS_G06214 | ERS11983154 | ERR9735524 | ERR9735524 | 1000000 |
| *Pterostylis tenuissima* Nicholls | *Foliosae* G.Don | *Diplodium tenuissimum* (Nicholls) D.L.Jones & M.A.Clem. | M.Duncan ORG 5146 (CANB 677104.1) | CNS_G06199 | ERS11983155 | ERR9735525 | ERR9735525 | 1000000 |
| *Pterostylis terminalis* (D.L.Jones & R.J.Bates) J.M.H.Shaw | *Oligochaetochilus* (Szlach.) Janes & Duretto | *Oligochaetochilus terminalis* D.L.Jones & R.J.Bates | M.A.Clements 11866 (CANB 950918.1) | CNS_G03417 | ERS11983156 | ERR9735526 | ERR9735526 | 0010000 |
| *Pterostylis torquata* D.L.Jones | *Foliosae* G.Don | *Diplodium torquatum* (D.L.Jones) D.L.Jones & M.A.Clem. | T.N.Hayashi 81 (CANB) | CNS_G05981 | ERS11983157 | ERR9735527 | ERR9735527 | 1000000 |
| *Pterostylis truncata* Fitzg. | *Foliosae* G.Don | *Diplodium truncatum* (Fitzg.) D.L.Jones & M.A.Clem. | G.Bradburn 13 (CANB) | CNS_G04996 | ERS11983158 | ERR9735528 | ERR9735528 | 1000000 |
| *Pterostylis tunstallii* D.L.Jones & M.A.Clem. | *Squamatae* G.Don | *Bunochilus tunstallii* (D.L.Jones & M.A.Clem.) D.L.Jones & M.A.Clem. | M.A.Clements 11802 (CANB) | CNS_G06203 | ERS11983159 | ERR9735529 | ERR9735529 | 1000000 |
| *Pterostylis turfosa* Endl. | *Catochilus* Benth. | *Plumatichilos turfosus* (Endl.) Szlach. | M.A.Clements 12309 & L. Nauheimer (CANB 950939.1) | CNS_G05949 | ERS11983160 | ERR9735530 | ERR9735530 | 0100000 |
| *Pterostylis valida* (Nicholls) D.L.Jones | *Oligochaetochilus* (Szlach.) Janes & Duretto | *Oligochaetochilus validus* (Nicholls) D.L.Jones & M.A.Clem. | J.Whitfield ORG6159 (CANB 755628.1) | CNS_G06213 | ERS11983161 | ERR9735531 | ERR9735531 | 1010000 |
| *Pterostylis venosa* Colenso | *Pterostylis* R.Br. | *Pterostylis venosa* Colenso | D.L.Jones s/n (CANB 787778.1) | CNS_G07751 | ERS11983162 | – | ERR9735532 | – |
| *Pterostylis ventricosa* (D.L.Jones) G.N.Backh. | *Parviflorae* (Bent.) Janes & Duretto | *Speculantha ventricosa* D.L.Jones | A.Stephenson ORG7309 (CANB) | CNS_G04340 | ERS11983163 | ERR9735533 | ERR9735533 | 1000000 |
| *Pterostylis vittata* Lindl. | *Urochilus* (D.L.Jones & M.A.Clem.) Janes & Duretto | *Urochilus vittatus* (Lindl.) D.L.Jones & M.A.Clem. | M.A.Clements 11939 (CANB 891174.1) | CNS_G01353 | ERS11983164 | ERR9735534 | ERR9735534 | 0100000 |
| *Pterostylis williamsonii* D.L.Jones | *Squamatae* G.Don | *Bunochilus williamsonii* (D.L.Jones) D.L.Jones & M.A.Clem. | M.A.Clements 10779A (CANB 891077.1) | CNS_G06218 | ERS11983165 | ERR9735535 | ERR9735535 | 1000000 |
| *Pterostylis xerophila* M.A.Clem. | *Oligochaetochilus* (Szlach.) Janes & Duretto | *Oligochaetochilus xerophilus* (M.A.Clem.) Szlach. | M.A.Clements 11595 (CANB 882126.1) | CNS_G06201 | ERS11983166 | ERR9735536 | ERR9735536 | 1010000 |
| **Outgroups** | | | | | | | | |
| *Adenochilus gracilis* Hook.f. | | *Adenochilus gracilis* Hook.f. | B.P.J.Molloy 174/00 (CHR 532785) | CNS_G04014 | ERS11983044 | ERR9735414 | ERR9735414 | – |
| *Anathallis obovata* (Lindl.) Pridgeon & M.W.Chase | | *Anathallis obovata* (Lindl.) Pridgeon & M.W.Chase | GenBank accession | | | NC043905* | – | – |
| *Aphyllorchis anomala* Dockrill | | *Aphyllorchis anomala* Dockrill | K.Schulte 143 (CNS 144311.1) | CNS_G00168 | ERS11983045 | ERR9735415 | – | – |
| *Apostasia wallichii* R.Br. | | *Apostasia stylidioides* (F.Muell.) Rchb.f. | K.Schulte 258 (CNS) | CNS_G06011 | ERS11983046 | ERR9735416 | – | – |
| *Calypso bulbosa* (L.) Oakes | | *Calypso bulbosa* (L.) Oakes | GenBank accession | | | NC040980* | – | – |
| *Cheirostylis notialis* D.L.Jones | | *Cheirostylis notialis* D.L.Jones | J.Moye ORG6978 (CANB 950940.1) | CNS_G03406 | ERS11983047 | ERR9735416 | – | – |
| *Chloraea gavilu* Lindl. | | *Chloraea gavilu* Lindl. | Givnish et al. 2018 | | | Givnish et al. 2018 | – | – |
| *Chloraea gavilu* Lindl. | | *Chloraea gavilu* Lindl. | GenBank accession | | | – | FR832121* | – |
| *Chloraea multiflora* Lindl. | | *Chloraea multiflora* Lindl. | GenBank accession | | | – | FR832119* | – |

| Species | Species *sensu* Jones & Clements[1] | Voucher details | DNA extraction no. | ENA sample accession number | Reference plastid data | Reference nuclear data | Area coding (abcdefg) |
|---|---|---|---|---|---|---|---|
| *Codonorchis lessonii* (d'Urv.) Lindl. | *Codonorchis lessonii* (d'Urv.) Lindl. | E.Pisano & O.Dollenz 5832 (CANB 950903.1) | CNS_G05327 | ERS11983048 | ERR9735418 | ERR9735418 | – |
| *Coilochilus neocaledonicum* Schltr. | *Coilochilus neocaledonicum* Schltr. | M.A.Clements 11248 (CANB 950932.1) | CNS_G05305 | ERS11983049 | ERR9735419 | – | – |
| *Cooktownia robertsii* D.L.Jones | *Cooktownia robertsii* D.L.Jones | L.J.Roberts (ORG 2113) (CANB 679460.1) | CNS_G06630 | ERS11983050 | ERR9735420 | – | – |
| *Cypripedium japonicum* Thunb. | *Cypripedium japonicum* Thunb. | GenBank accession | | | NC027227* | – | – |
| *Danhatchia australis* (Hatch) Garay & Christenson | *Danhatchia novaehollandiae* D.L.Jones & M.A.Clem. | P.H.Weston, W.A.Cherry & J. Stockard 3405 (NSW 870437) | CNS_G03415 | ERS11983051 | ERR9735421 | – | – |
| *Dendrobium aduncum* Lindl. | *Dendrobium aduncum* Lindl. | GenBank accession | | | NC038077* | – | – |
| *Dendrobium flexicaule* Z.H.Tsi, S.C.Sun & L.G.Xu | *Dendrobium flexicaule* Z.H.Tsi, S.C.Sun & L.G.Xu | GenBank accession | | | NC038076* | – | – |
| *Disa bracteata* Sw. | *Disa bracteata* Sw. | M.A.Clements & L.Nauheimer 12256 (CANB 906207.1) | CNS_G05322 | ERS11983052 | ERR9735422 | ERR9735422 | – |
| *Diuris sulphurea* R.Br. | *Diuris sulphurea* R.Br. | M.A.Clements 12053 (CANB 908278.1) | CNS_G02433 | ERS11983053 | ERR9735423 | ERR9735423 | – |
| *Elleanthus sodiroi* Schltr. | *Elleanthus sodiroi* Schltr. | GenBank accession | | | NC027266* | – | – |
| *Epipactis palustris* (L.) Crantz | *Epipactis palustris* (L.) Crantz | GenBank accession | | | NC041187* | – | – |
| *Eucosia umbrosa* D.L.Jones & M.A.Clem. | *Eucosia umbrosa* D.L.Jones & M.A.Clem. | D.L.Jones s.n. (CANB 676421.1) | CNS_G06632 | ERS11983054 | ERR9735424 | – | – |
| *Eucosia viridiflora* (Blume) M.C.Pace | *Goodyera viridiflora* (Blume) Blume | W.K.Harris 224 (CANB 950917.1) | CNS_G01321 | ERS11983055 | ERR9735425 | – | – |
| *Eulophia graminea* Lindl. | *Eulophia graminea* Lindl. | C.P.Brock 311 (CANB 596921.1) | CNS_G02766 | ERS11983056 | ERR9735426 | – | – |
| *Gastrochilus calceolaris* (Buch.-Ham. ex Sm.) D.Don | *Gastrochilus calceolaris* (Buch.-Ham. ex Sm.) D.Don | GenBank accession | | | NC042686* | – | – |
| *Goodyera schlechtendaliana* Rchb.f. | *Goodyera schlechtendaliana* Rchb.f. | M.A.Clements 12186 (CANB 950929.1) | CNS_G04898 | ERS11983057 | ERR9735427 | ERR9735427 | – |
| *Habenaria elongata* R.Br. | *Pecteilis elongata* R.Br. | A.R.Field 3904 (CNS) | CNS_G03456 | ERS11983058 | ERR9735428 | ERR9735428 | – |
| *Hetaeria oblongifolia* Blume | *Hetaeria oblongifolia* Blume | K.Schulte 150 (CNS 144275.1) | CNS_G00729 | ERS11983059 | ERR9735429 | – | – |
| *Holcoglossum subulifolium* (Rchb.f.) Christenson | *Holcoglossum subulifolium* (Rchb.f.) Christenson | GenBank accession | | | NC041519* | – | – |
| *Neottia japonica* (Blume) Szlach. | *Neottia japonica* (Blume) Szlach. | GenBank accession | | | NC041446* | – | – |
| *Nervilia aragoana* Gaudich. | *Nervilia aragoana* Gaudich. | L.Roberts ORG3787 (CANB 669943.1) | CNS_G03442 | ERS11983060 | ERR9735430 | ERR9735430 | – |
| *Paphiopedilum delenatii* Guillaumin | *Paphiopedilum delenatii* Guillaumin | GenBank accession | | | NC041309* | – | – |
| *Pendulorchis himalaica* (Deb, Sengupta & Malick) Z.J.Liu, K.Wei Liu & X.J.Xiao | *Pendulorchis himalaica* (Deb, Sengupta & Malick) Z.J.Liu, K.Wei Liu & X.J.Xiao | GenBank accession | | | NC041513* | – | – |
| *Peristylus banfieldii* (F.M.Bailey) Lavarack | *Peristylus banfieldii* (F.M.Bailey) Lavarack | R.Collins 1158 (CANB 7904962.1) | CNS_G06628 | ERS11983061 | ERR9735431 | – | – |
| *Phragmipedium longifolium* (Rchb.f. & Warsz.) Rolfe | *Phragmipedium longifolium* (Rchb.f. & Warsz.) Rolfe | GenBank accession | | | NC028149* | – | – |

| **Species** | **Species *sensu* Jones & Clements**[1] | **Voucher details** | **DNA extraction no.** | **ENA sample accession number** | **Reference plastid data** | **Reference nuclear data** | **Area coding (abcdefg)** |
|---|---|---|---|---|---|---|---|
| *Rhomboda polygonoides* (F.Muell.) Ormerod | *Rhomboda polygonoides* (F.Muell.) Ormerod | K.Schulte 249 (CNS 144274.1) | CNS_G00708 | ERS11983167 | ERR9735537 | – | – |
| *Salacistis rubicunda* (Blume) M.C.Pace | *Salacistis ochroleuca* (F.M.Bailey) M.A.Clem. & D.L.Jones | B.Gray 8333 (CANB 507310.1) | CNS_G06635 | ERS11983168 | ERR9735538 | – | – |
| *Sobralia callosa* L.O.Williams | *Sobralia callosa* L.O.Williams | GenBank accession | | | NC028147* | – | – |
| *Spiranthes aestivalis* (Poir.) Rich. | *Spiranthes aestivalis* (Poir.) Rich. | M.A.Clements 12199 (CANB 909010.1) | CNS_G05319 | ERS11983169 | ERR9735539 | ERR9735539 | – |
| *Spiranthes sinensis* (Pers.) Ames | *Spiranthes australis* Lindl. | T.N.Hayashi 67 | CNS_G04983 | ERS11983170 | ERR9735540 | ERR9735540 | – |
| *Townsonia viridis* (Hook.f.) Schltr. | *Townsonia viridis* (Hook.f.) Schltr. | H.Wapstra ORG 4598 (CANB 670901.1) | CNS_G01314 | ERS11983171 | ERR9735541 | – | – |
| *Vanilla aphylla* Blume | *Vanilla aphylla* Blume | GenBank accession | | | NC035320* | – | – |
| *Vanilla pompona* Schiede | *Vanilla pompona* Schiede | GenBank accession | | | NC036809* | – | – |
| *Zeuxine oblonga* R.S.Rogers & C.T.White | *Zeuxine oblonga* R.S.Rogers & C.T.White | K.Schulte 181 (CNS 144329.1) | CNS_G00175 | ERS11983172 | ERR9735542 | ERR9735542 | – |
