## Supplementary Material S2 for "Evolutionary relationships and range evolution of greenhood orchids (subtribe Pterostylidinae): insights from plastid phylogenomics"

Phylogenetic relationships in Pterostylidinae based on plastid and nuclear data.

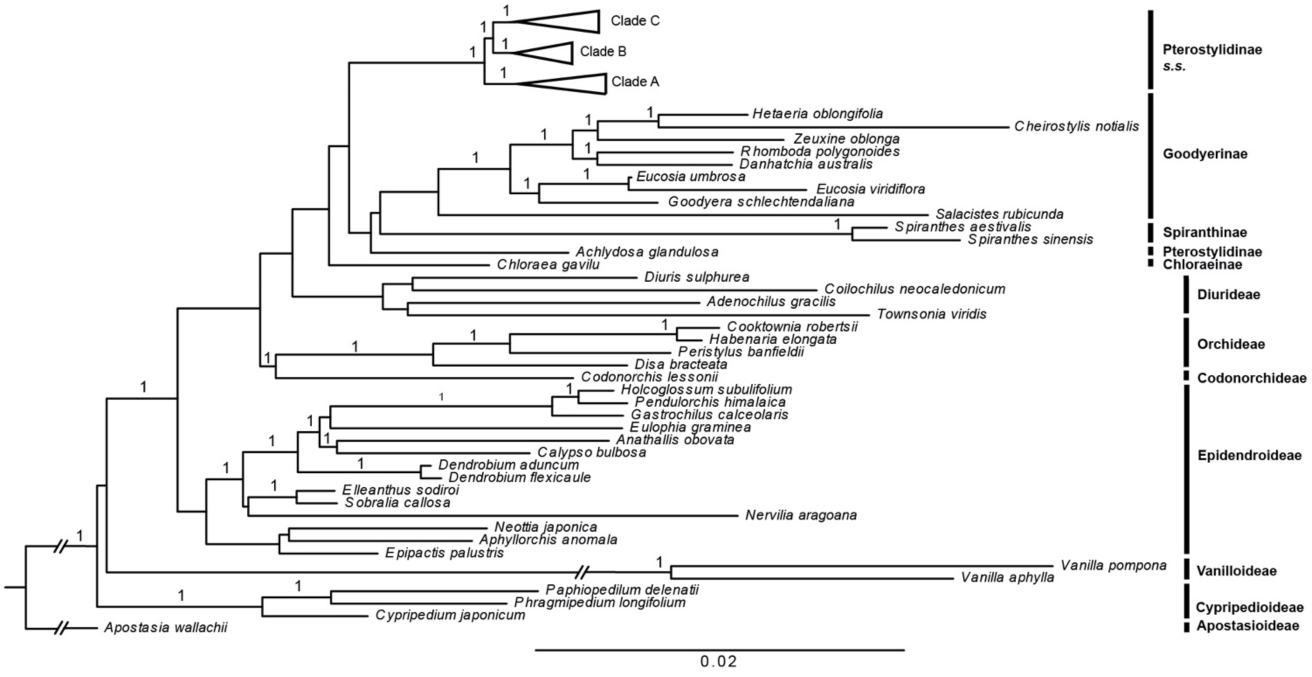


**Supplementary Material S2.1** Phylogenetic placement of Pterostylidinae within Orchidaceae. Bayesian inference based on 75 plastid genes (91,090 bp alignment). Nodal support values are given above branches (posterior probabilities > 0.95).

**Next page: Supplementary Material S2.2** Phylogenetic relationships in Pterostylidinae. Bayesian inference based on 75 plastid genes (91,090 bp alignment). Labels A, B, and C refer to the three major clades within the genus, C1 and C2 denote two main clades within Clade C. Nodal support values are given above branches (posterior probabilities > 0.95).

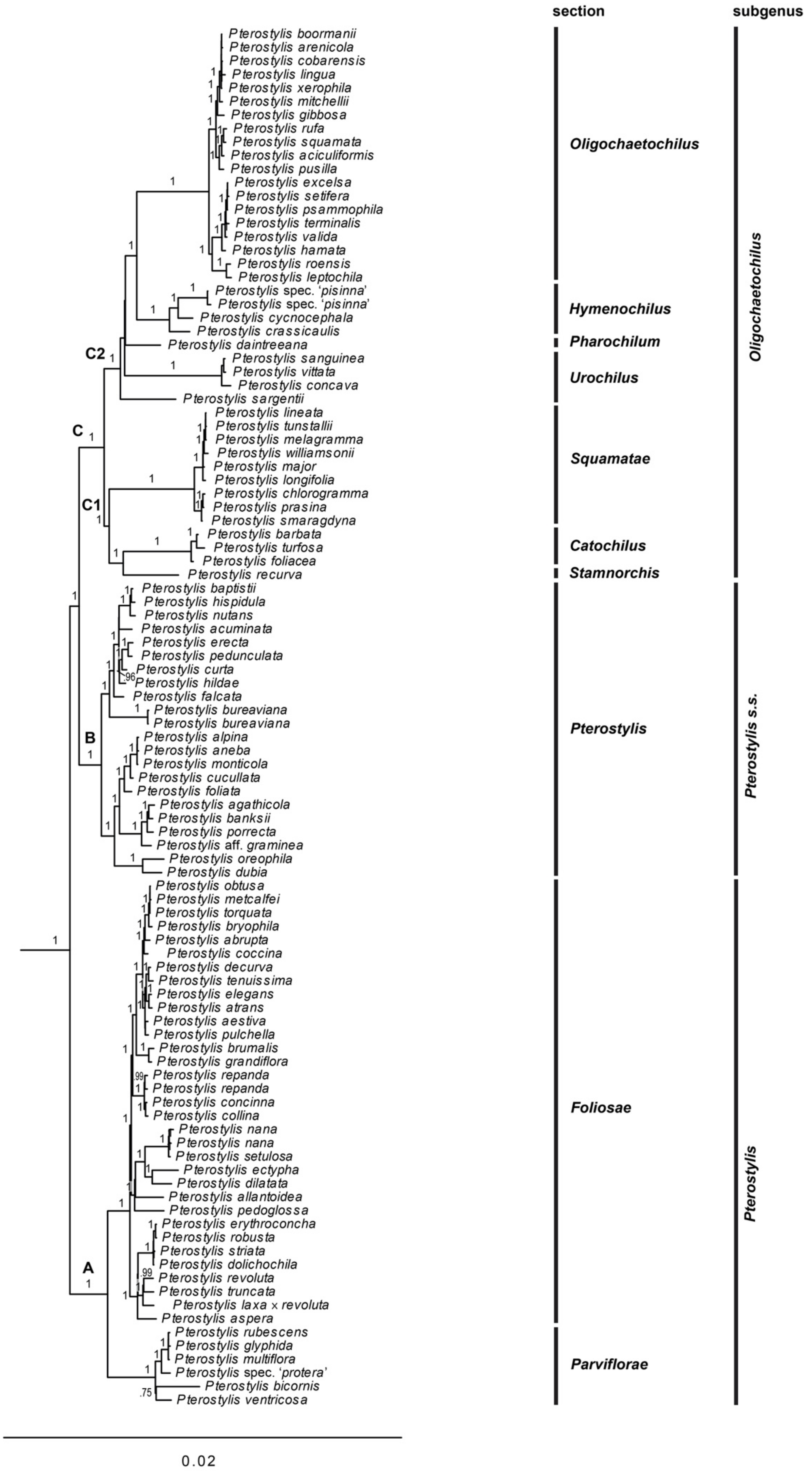
section
subgenus
Pterostylis boormanii
Pterostylis arenicola
Pterostylis cobarensis
Pterostylis lingua
Pterostylis xerophila
Pterostylis mitchellii
Pterostylis gibbosa
Pterostylis rufa
Pterostylis squamata
Pterostylis aciculiformis
Pterostylis pusilla
Pterostylis excelsa
Pterostylis setifera
Pterostylis psammophila
Pterostylis terminalis
Pterostylis valida
Pterostylis hamata
Pterostylis roensis
Pterostylis leptochila
Pterostylis spec. 'pisinna'
Pterostylis spec. 'pisinna'
Pterostylis cycnocephala
Pterostylis crassicaulis
Pterostylis daintreeana
Pterostylis sanguinea
Pterostylis vittata
Pterostylis concava
Pterostylis sargentii
Pterostylis lineata
Pterostylis tunstallii
Pterostylis melagramma
Pterostylis williamsonii
Pterostylis major
Pterostylis longifolia
Pterostylis chlorogramma
Pterostylis prasina
Pterostylis smaragdyna
Pterostylis barbata
Pterostylis turfosa
Pterostylis foliacea
Pterostylis recurva
Pterostylis baptistii
Pterostylis hispidula
Pterostylis nutans
Pterostylis acuminata
Pterostylis erecta
Pterostylis pedunculata
Pterostylis curta
Pterostylis hildae
Pterostylis falcata
Pterostylis bureaviana
Pterostylis bureaviana
Pterostylis alpina
Pterostylis aneba
Pterostylis monticola
Pterostylis cucullata
Pterostylis foliata
Pterostylis agathicola
Pterostylis banksii
Pterostylis porrecta
Pterostylis aff. graminea
Pterostylis oreophila
Pterostylis dubia
Pterostylis obtusa
Pterostylis metcalfei
Pterostylis torquata
Pterostylis bryophila
Pterostylis abrupta
Pterostylis coccina
Pterostylis decurva
Pterostylis tenuissima
Pterostylis elegans
Pterostylis atrans
Pterostylis aestiva
Pterostylis pulchella
Pterostylis brumalis
Pterostylis grandiflora
Pterostylis repanda
Pterostylis repanda
Pterostylis concinna
Pterostylis collina
Pterostylis nana
Pterostylis nana
Pterostylis setulosa
Pterostylis ectypha
Pterostylis dilatata
Pterostylis allantoidea
Pterostylis pedoglossa
Pterostylis erythroconcha
Pterostylis robusta
Pterostylis striata
Pterostylis dolichochila
Pterostylis revoluta
Pterostylis truncata
Pterostylis laxa × revoluta
Pterostylis aspera
Pterostylis rubescens
Pterostylis glyphida
Pterostylis multiflora
Pterostylis spec. 'protera'
Pterostylis bicornis
Pterostylis ventricosa
C2
C
C1
B
A
Oligochaetochilus
Hymenochilus
Pharochilum
Urochilus
Squamatae
Catochilus
Stamnorchis
Pterostylis
Foliosae
Parviflorae
Oligochaetochilus
Pterostylis s.s.
Pterostylis
0.02

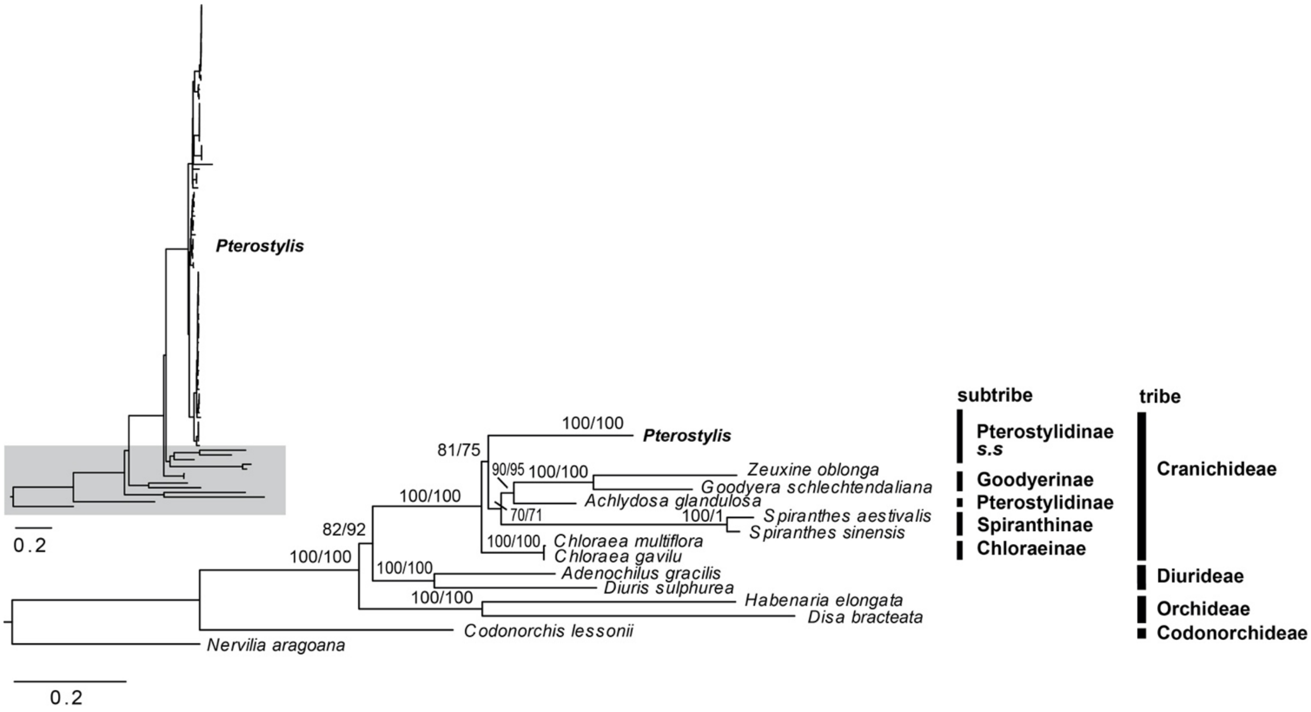


**Supplementary Material S2.3** Phylogenetic position of Pterostylidinae within Orchidoideae. Maximum likelihood (ML) reconstruction in RaxML based on the ribosomal RNA cistron (8,808 bp alignment). Nodal support values are given above branches (bootstrap values above 50 from RAxML analysis, followed by ultrafast bootstrap values from IQTREE analysis).

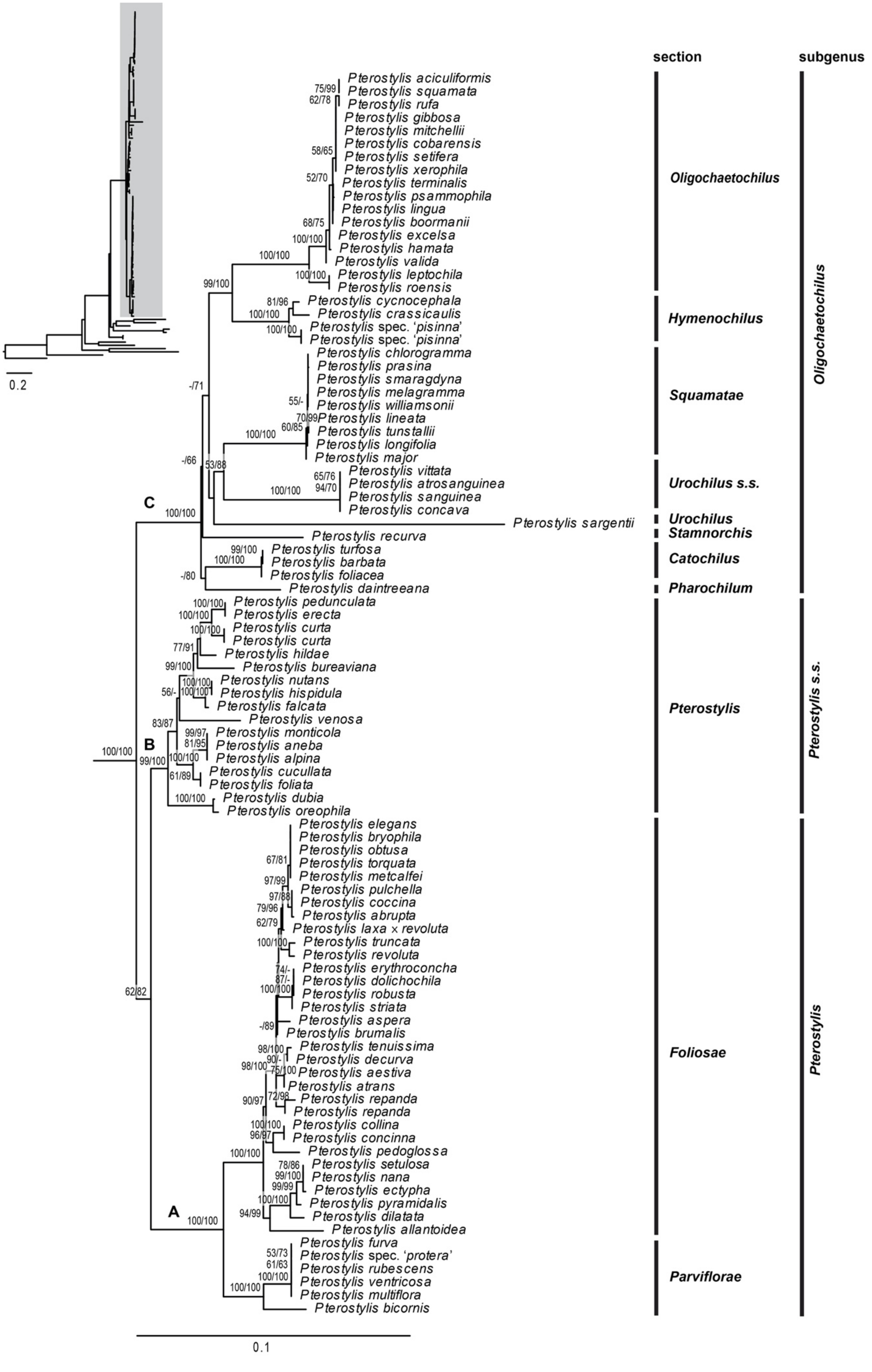
0.2
section
subgenus
Pterostylis aciculiformis
Pterostylis squamata
Pterostylis rufa
Pterostylis gibbosa
Pterostylis mitchellii
Pterostylis cobarensis
Pterostylis setifera
Pterostylis xerophila
Pterostylis terminalis
Pterostylis psammophila
Pterostylis lingua
Pterostylis boormanii
Pterostylis excelsa
Pterostylis hamata
Pterostylis valida
Pterostylis leptochila
Pterostylis roensis
Pterostylis cycnocephala
Pterostylis crassicaulis
Pterostylis spec. 'pisinna'
Pterostylis spec. 'pisinna'
Pterostylis chlorogramma
Pterostylis prasina
Pterostylis smaragdyna
Pterostylis melagramma
Pterostylis williamsonii
Pterostylis lineata
Pterostylis tunstallii
Pterostylis longifolia
Pterostylis major
Pterostylis vittata
Pterostylis atrosanguinea
Pterostylis sanguinea
Pterostylis concava
Pterostylis sargentii
Pterostylis recurva
Pterostylis turfosa
Pterostylis barbata
Pterostylis foliacea
Pterostylis daintreeana
Pterostylis pedunculata
Pterostylis erecta
Pterostylis curta
Pterostylis curta
Pterostylis hildae
Pterostylis bureaviana
Pterostylis nutans
Pterostylis hispidula
Pterostylis falcata
Pterostylis venosa
Pterostylis monticola
Pterostylis aneba
Pterostylis alpina
Pterostylis cucullata
Pterostylis foliata
Pterostylis dubia
Pterostylis oreophila
Pterostylis elegans
Pterostylis bryophila
Pterostylis obtusa
Pterostylis torquata
Pterostylis metcalfei
Pterostylis pulchella
Pterostylis coccina
Pterostylis abrupta
Pterostylis laxa × revoluta
Pterostylis truncata
Pterostylis revoluta
Pterostylis erythroconcha
Pterostylis dolichochila
Pterostylis robusta
Pterostylis striata
Pterostylis aspera
Pterostylis brumalis
Pterostylis tenuissima
Pterostylis decurva
Pterostylis aestiva
Pterostylis atrans
Pterostylis repanda
Pterostylis repanda
Pterostylis collina
Pterostylis concinna
Pterostylis pedoglossa
Pterostylis setulosa
Pterostylis nana
Pterostylis ectypha
Pterostylis pyramidalis
Pterostylis dilatata
Pterostylis allantoidea
Pterostylis furva
Pterostylis spec. 'protera'
Pterostylis rubescens
Pterostylis ventricosa
Pterostylis multiflora
Pterostylis bicornis
A
B
C
Oligochaetochilus
Hymenochilus
Squamatae
Urochilus s.s.
Urochilus
Stamnorchis
Catochilus
Pharochilum
Pterostylis
Foliosae
Parviflorae
Oligochaetochilus
Pterostylis s.s.
Pterostylis
0.1

**Previous page: Supplementary Material S2.4:** Phylogenetic relationships in Pterostylidinae. Maximum likelihood (ML) reconstruction in RaxML based on the ribosomal RNA cistron (8,808 bp alignment). Labels A, B, and C refer to the three major clades within the genus, C1 and C2 denote two main clades within Clade C. Nodal support values are given above branches (bootstrap values > 50 from RAxML analysis, followed by ultrafast bootstrap values from IQTREE analysis).

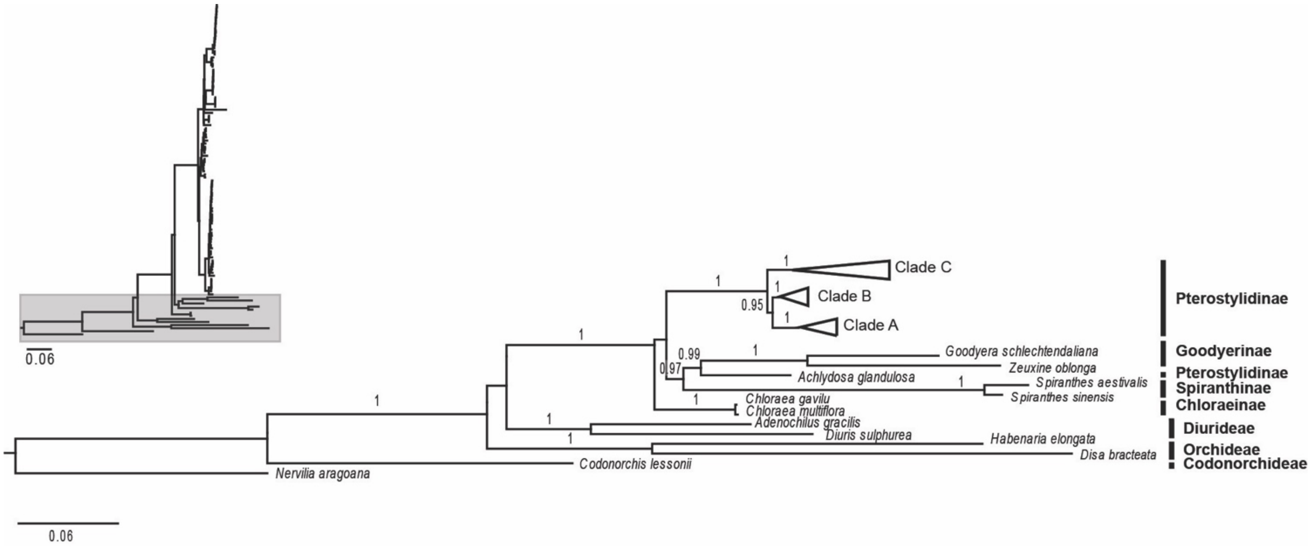


**Supplementary Material S2.5**. Phylogenetic placement of Pterostylidinae in Orchidoideae. Bayesian inference based on the ribosomal RNA cistron (8,808 bp alignment). Nodal support values are given above branches (posterior probabilities > 0.95).

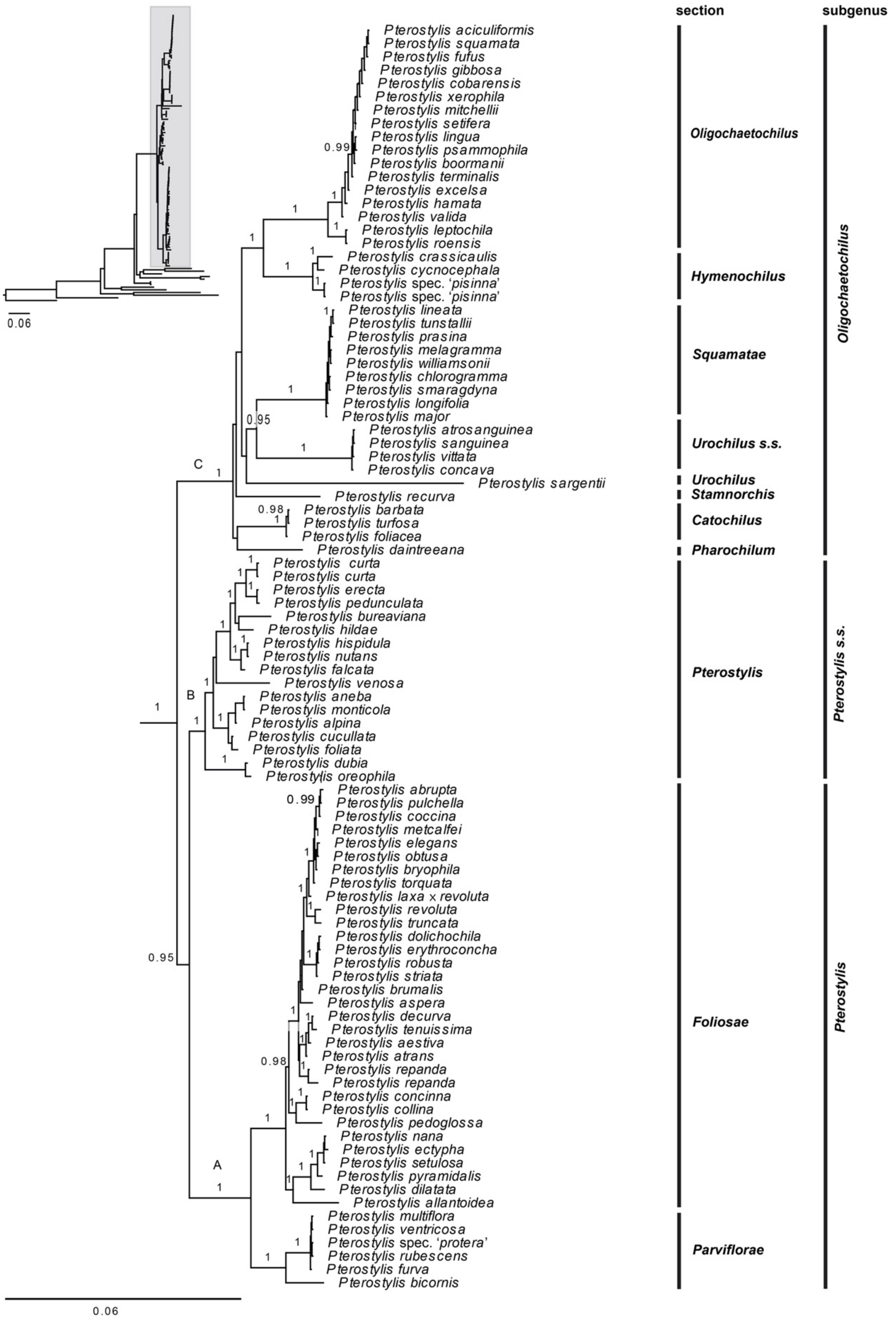


**Supplementary Material S2.6**. Phylogenetic relationships in Pterostylidinae. Bayesian inference based on the ribosomal RNA cistron (8,808 bp alignment). Labels A, B, and C refer to the three major clades within the genus. Nodal support values are given above branches (posterior probabilities > 0.95).
