## Supplementary Material S3 for "Evolutionary relationships and range evolution of greenhood orchids (subtribe Pterostylidinae): insights from plastid phylogenomics"

**Divergence time estimations based on plastid and nuclear data.**

**Supplementary Material 3.1.** Selection of the best fit models for divergence time estimation based on the 25 most parsimony informative plastid genes. Molecular clock and speciation models were evaluated using a posterior simulation-based analogue of Akaike's information criterion (AIC; Akaike, 1974), termed AICM (Raftery et al., 2007 as implemented in AICM Analyser included in BEASTv2.6.1. The best fit model for each dataset is highlighted in bold. The most parsimony informative plastid genes in descending order: *rpo*C2, *ycf*2, *rpo*B, *mat*K, *acc*D, *ndh*F, *rpo*C1, *ccs*A, *psa*B, *rps*16, *psa*A, *atp*A, *ndh*D, *rpo*A, *atp*B, *psb*B, *clp*P, *rps*2, *cem*A, *rps*3, *ndh*A, *psb*C, *atp*F, and *psb*A.

| Dataset | Clock model | Model for speciation/extinction process | AICM |
|---|---|---|---|
| **Plastid** | pure birth (Yule) | uncorrelated relaxed lognormal clock | 418029.1158 |
| | pure birth (Yule) | strict clock | 422354.0833 |
| | birth death | uncorrelated relaxed lognormal clock | **418019.8344** |
| | birth death | strict clock | 422336.9326 |
| **Nuclear** | pure birth (Yule) | uncorrelated relaxed lognormal clock | **89144.3379** |
| | pure birth (Yule) | strict clock | 89759.142 |
| | birth death | uncorrelated relaxed lognormal clock | 89161.9538 |
| | birth death | strict clock | 89689.3996 |

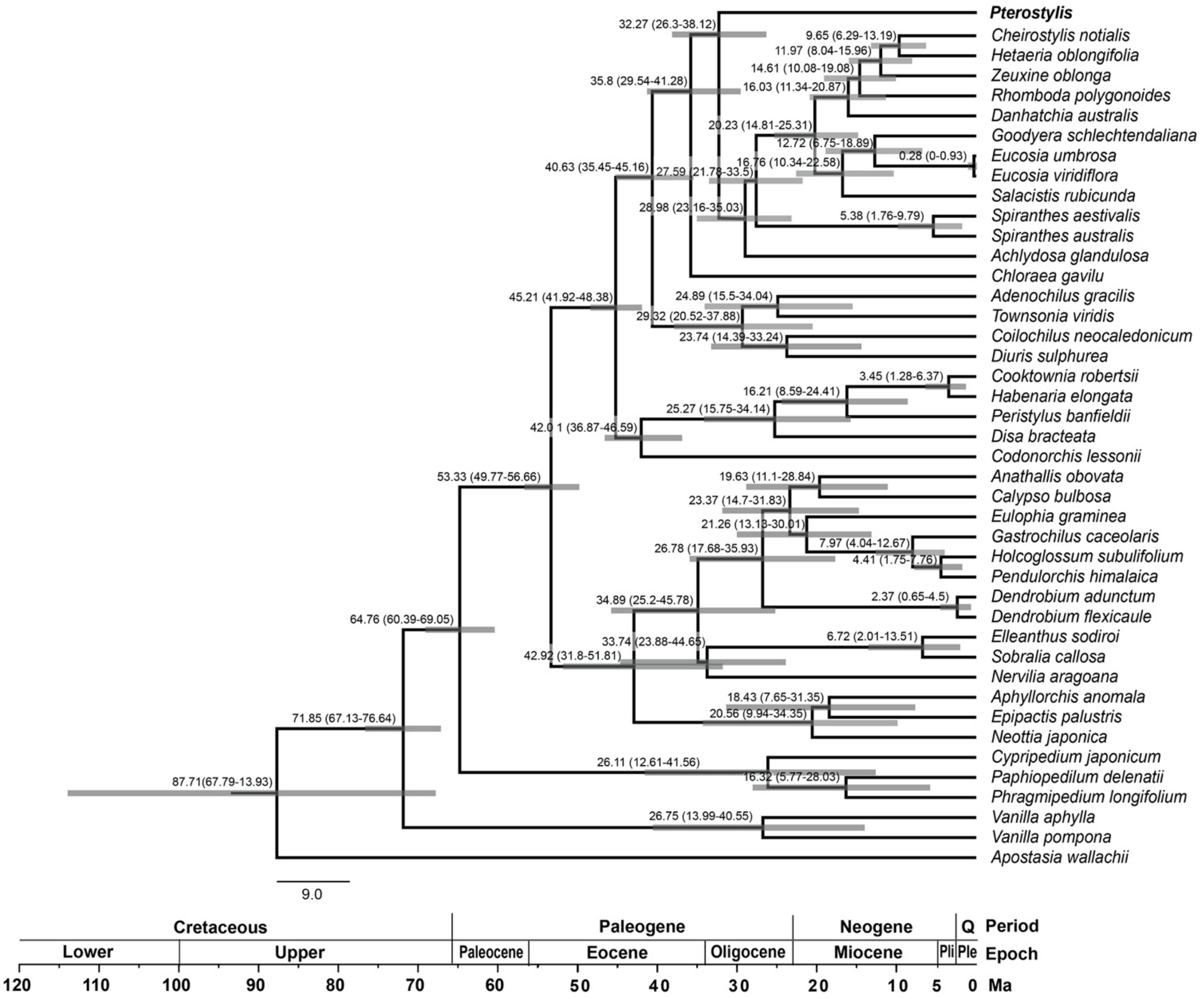


**Supplementary Material 3.2.** Chronogram showing divergence time estimations within Orchidaceae. Maximum clade credibility tree from Bayesian divergence time estimation based on 25 most informative plastid genes and an uncorrelated molecular clock model under the birth-death tree prior. Divergence times (Ma) are given at each node together with 95% highest posterior density (HDP) values indicated by grey bars and numbers in brackets.

**Next page: Supplementary Material 3.3.** Chronogram showing divergence times within Pterostylidinae *s.s.*. based on plastid data. Maximum clade credibility tree from Bayesian divergence time estimation based on 25 most informative plastid genes and an uncorrelated molecular clock model under the birth-death tree prior. Divergence times (Ma) are given at each node together with 95% highest posterior density (HDP) values indicated by grey bars and numbers in brackets.

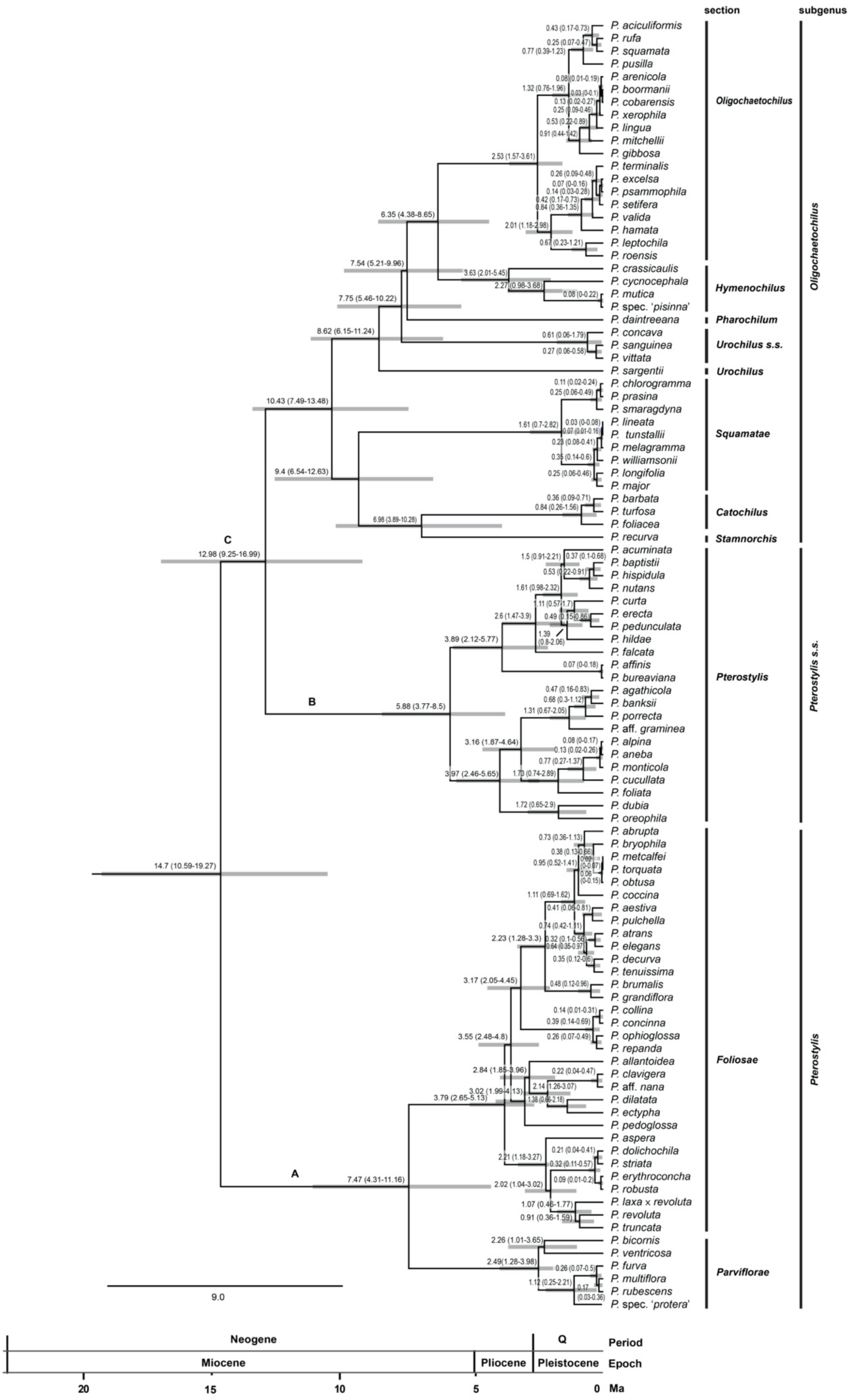

section
subgenus
P. aciculiformis
P. rufa
P. squamata
P. pusilla
P. arenicola
P. boormanii
P. cobarensis
P. xerophila
P. lingua
P. mitchellii
P. gibbosa
P. terminalis
P. excelsa
P. psammophila
P. setifera
P. valida
P. hamata
P. leptochila
P. roensis
P. crassicaulis
P. cycnocephala
P. mutica
P. spec. 'pisinna'
P. daintreeana
P. concava
P. sanguinea
P. vittata
P. sargentii
P. chlorogramma
P. prasina
P. smaragdyna
P. lineata
P. tunstallii
P. melagramma
P. williamsonii
P. longifolia
P. major
P. barbata
P. turfosa
P. foliacea
P. recurva
P. acuminata
P. baptistii
P. hispidula
P. nutans
P. curta
P. erecta
P. pedunculata
P. hildae
P. falcata
P. affinis
P. bureaviana
P. agathicola
P. banksii
P. porrecta
P. aff. graminea
P. alpina
P. aneba
P. monticola
P. cucullata
P. foliata
P. dubia
P. oreophila
P. abrupta
P. bryophila
P. metcalfei
P. torquata
P. obtusa
P. coccina
P. aestiva
P. pulchella
P. atrans
P. elegans
P. decurva
P. tenuissima
P. brumalis
P. grandiflora
P. collina
P. concinna
P. ophioglossa
P. repanda
P. allantoidea
P. clavigera
P. aff. nana
P. dilatata
P. ectypha
P. pedoglossa
P. aspera
P. dolichochila
P. striata
P. erythroconcha
P. robusta
P. laxa × revoluta
P. revoluta
P. truncata
P. bicornis
P. ventricosa
P. furva
P. multiflora
P. rubescens
P. spec. 'protera'
Oligochaetochilus
Hymenochilus
Pharochilum
Urochilus s.s.
Urochilus
Squamatae
Catochilus
Stamnorchis
Pterostylis
Foliosae
Parviflorae
Oligochaetochilus
Pterostylis s.s.
Pterostylis
C
B
A
14.7 (10.59-19.27)
12.98 (9.25-16.99)
10.43 (7.49-13.48)
9.4 (6.54-12.63)
8.62 (6.15-11.24)
7.75 (5.46-10.22)
7.54 (5.21-9.96)
6.35 (4.38-8.65)
5.88 (3.77-8.5)
7.47 (4.31-11.16)
3.79 (2.65-5.13)
9.0
Neogene
Q
Period
Miocene
Pliocene
Pleistocene
Epoch
20
15
10
5
0
Ma

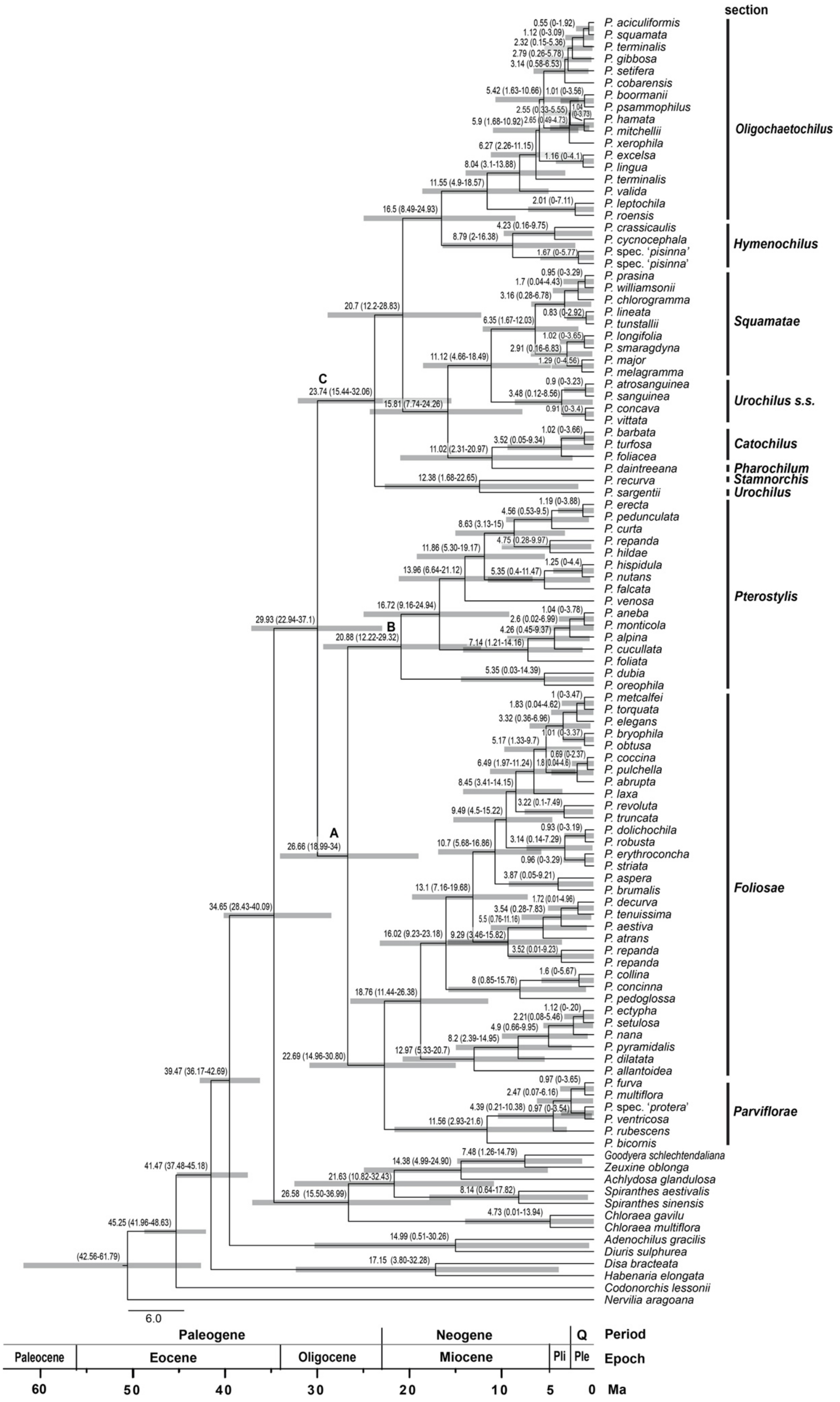
section
P. aciculiformis
P. squamata
P. terminalis
P. gibbosa
P. setifera
P. cobarensis
P. boormanii
P. psammophilus
P. hamata
P. mitchellii
P. xerophila
P. excelsa
P. lingua
P. terminalis
P. valida
P. leptochila
P. roensis
Oligochaetochilus
P. crassicaulis
P. cycnocephala
P. spec. 'pisinna'
P. spec. 'pisinna'
Hymenochilus
P. prasina
P. williamsonii
P. chlorogramma
P. lineata
P. tunstallii
P. longifolia
P. smaragdyna
P. major
P. melagramma
Squamatae
P. atrosanguinea
P. sanguinea
P. concava
P. vittata
Urochilus s.s.
P. barbata
P. turfosa
P. foliacea
Catochilus
P. daintreeana
Pharochilum
P. recurva
Stamnorchis
P. sargentii
Urochilus
P. erecta
P. pedunculata
P. curta
P. repanda
P. hildae
P. hispidula
P. nutans
P. falcata
P. venosa
P. aneba
P. monticola
P. alpina
P. cucullata
P. foliata
P. dubia
P. oreophila
Pterostylis
P. metcalfei
P. torquata
P. elegans
P. bryophila
P. obtusa
P. coccina
P. pulchella
P. abrupta
P. laxa
P. revoluta
P. truncata
P. dolichochila
P. robusta
P. erythroconcha
P. striata
P. aspera
P. brumalis
P. decurva
P. tenuissima
P. aestiva
P. atrans
P. repanda
P. repanda
P. collina
P. concinna
P. pedoglossa
P. ectypha
P. setulosa
P. nana
P. pyramidalis
P. dilatata
P. allantoidea
Foliosae
P. furva
P. multiflora
P. spec. 'protera'
P. ventricosa
P. rubescens
P. bicornis
Parviflorae
Goodyera schlechtendaliana
Zeuxine oblonga
Achlydosa glandulosa
Spiranthes aestivalis
Spiranthes sinensis
Chloraea gavilu
Chloraea multiflora
Adenochilus gracilis
Diuris sulphurea
Disa bracteata
Habenaria elongata
Codonorchis lessonii
Nervilia aragoana
A
B
C
6.0
Paleogene
Neogene
Q
Period
Paleocene
Eocene
Oligocene
Miocene
Pli
Ple
Epoch
60
50
40
30
20
10
5
0
Ma

**Previous page: Supplementary Material S3.4.** Chronogram showing divergence times within Pterostylidinae *s.s.* based on nuclear data. Maximum clade credibility tree from Bayesian divergence time estimation based on nuclear ribosomal RNA cistron and an uncorrelated molecular clock model under the Yule tree prior. Divergence times (Ma) are given at each node together with 95% highest posterior density (HDP) values indicated by grey bars and numbers in brackets.

**Supplementary Material S3.5.** Comparison of divergence time estimations of main lineages within Pterostylidinae *s.s.* derived from plastid and nuclear data. HDP: 95% highest posterior density.

| | **Divergence time estimate based on plastid data** | | | **Divergence time estimate based on nuclear data** | | |
|---|---|---|---|---|---|---|
| **Node** | **Ma** | **HDP range** | **epoch** | **Ma** | **HDP range** | **epoch** |
| Pterostylidinae s.s. stem | 32.27 | 26.3-38.12 | Oligocene | 34.65 | 28.43-40.09 | Oligocene |
| Pterostylidinae s.s. crown | 14.7 | 10.59-19.27 | mid Miocene | 29.93 | 22.94-37.1 | early Oligocene |
| *Pterostylis* Clade A crown | 7.47 | 4.31-11.16 | late Miocene | 22.69 | 14.96-30.90 | early Miocene |
| Pterostylis Clade B crown | 5.88 | 3.77-8.5 | late Miocene | 20.88 | 12.22-29.32 | early Miocene |
| Pterostylis Clade C crown | 10.43 | 7.49-13.48 | late Miocene | 23.74 | 15.44-32.06 | late Oligocene |
| *P.* sect. *Speculantha* stem | 7.47 | 4.31-11.16 | late Miocene | 22.69 | 14.96-30.80 | early Miocene |
| *P.* sect. *Speculantha* crown | 2.49 | 1.28-3.98 | early Pleistocene | 11.56 | 2.93-21.6 | late Miocene |
| *P.* sect. *Foliosae* stem | 7.47 | 4.31-11.16 | late Miocene | 22.69 | 14.96-30.8 | early Miocene |
| *P.* sect. *Foliosae* crown | 3.79 | 2.65-5.13 | early Pliocene | 18.76 | 11.44-26.38 | early Miocene |
| *P.* sect. *Pterostylis* stem | 12.98 | 9.25-16.99 | mid Miocene | 29.93 | 22.94-37.1 | early Oligocene |
| *P.* sect. *Pterostylis* crown | 5.88 | 3.77-8.5 | late Miocene | 20.88 | 12.22-29.32 | mid Miocene |
| *P.* sect. *Stamnorchis* stem | 6.98 | 3.89-10.28 | late Miocene | 12.38 | 1.68-22.68 | mid Miocene |
| *P.* sect. *Catochilus* stem | 6.98 | 3.89-10.28 | late Miocene | 15.81 | 7.74-24.26 | early Miocene |
| *P.* sect. *Catochilus* crown | 0.84 | 0.26-1.56 | early Pleistocene | 10.02 | 2.31-20.97 | late Miocene |
| *P.* sect. *Urochilus* s.s. stem | 7.75 | 5.46-10.22 | late Miocene | 11.12 | 4.66-18.49 | late Miocene |
| *P.* sect. *Urochilus* s.s. crown | 0.61 | 0.06-1.79 | mid Pleistocene | 3.48 | 0.12-8.56 | late Pleiocene |
| *P.* sect. *Squamatae* stem | 9.4 | 6.54-12.63 | late Miocene | 11.12 | 4.66-18.49 | late Miocene |
| *P.* sect. *Squamatae* crown | 1.61 | 0.7-2.82 | early Pleistocene | 6.35 | 1.67-12.03 | late Miocene |
| *P.* sect. *Hymenochilus* stem | 6.35 | 4.38-8.65 | late Miocene | 16.5 | 8.49-24.93 | early Miocene |
| *P.* sect. *Hymenochilus* crown | 3.63 | 2.01-5.45 | early Pliocene | 8.79 | 2-16.38 | late Miocene |
| *P.* sect. *Oligochaetochilus* stem | 6.35 | 4.38-8.65 | late Miocene | 16.5 | 8.49-24.93 | early Miocene |
| *P.* sect. *Oligochaetochilus* crown | 2.53 | 1.57-3.61 | late Pliocene | 11.55 | 4.9-18.57 | mid Miocene |
| *P. sargentii* stem | 8.62 | 6.15-11.24 | late Miocene | 12.38 | 1.68-22.65 | mid Miocene |
